# Structural basis for auto-inhibition and activation of a short prokaryotic Argonaute associated TIR-APAZ defense system

**DOI:** 10.1101/2023.07.12.548734

**Authors:** Lijie Guo, Pingping Huang, Zhaoxin Li, Young-Cheul Shin, Purui Yan, Meiling Lu, Meirong Chen, Yibei Xiao

## Abstract

Short prokaryotic Ago accounts for most prokaryotic Argonaute (pAgo) and is involved in defending bacteria against invading nucleic acids. Short prokaryotic Ago associated with APAZ-TIR (SPARTA) has been shown to oligomerize and deplete NAD^+^ upon guide-mediated target DNA recognition. However, the molecular basis of SPARTA inhibition and activation remains unknown. Here we determine the cryo-EM structures of *Crenotalea thermophila* SPARTA in its inhibited, transient, as well as activated states. The SPARTA is auto-inhibited by its acidic tail, which occupies the guide-target binding channel. Guide mediated target binding expels this acidic tail and triggers substantial conformational changes to expose Ago-Ago dimerization interface. As a result, SPARTA assembles into an active TIR-APAZ_4_/short Ago_4_ octamer, where the four TIR domains are rearranged and packed to form NADase active sites. Together with biochemical evidence, our results provide a panoramic vision explaining SPARTA auto-inhibition and activation, and expand understanding of pAgo mediated bacterial defense systems.

## Introduction

Argonaute proteins (Agos) are defense elements found in both eukaryotes and prokaryotes, utilizing small (typically 15–30 nucleotides long) oligonucleotides as guides to target complementary nucleic acids. All eukaryotic Ago (eAgo) possess an N (N-terminal) domain, a PAZ (PIWI-Argonaute-Zwille) domain, a MID (middle) domain, and a PIWI (P-element Induced Wimpy Testis) domain constituting a bi-lobed scaffold to employ ssRNA-guided complementary RNA silencing ^[1–3]^. Prokaryotic Ago (pAgo) show much higher diversity than eAgo, and can be generally classified into long-A Ago, long-B Ago and short Ago ^[4–7]^. The short Ago accounts for over half of pAgo but lacks the N domain and PAZ domain ^[4]^. This “imperfect” short Ago is normally complemented by an APAZ (analog of PAZ) domain encoded in the same operon, which is mostly appended with another functional domain, including Toll-interleukin receptor (TIR), Silent information regulator 2 (SIR2), Schlafen/Alba, Mrr-like, RecB, and RecG/DHS-like domain ^[4,6,7,9]^. Among them, the APAZ/short Ago fused with TIR and Sir2 are most abundant and have been implicated in host defense ^[10–12]^. However, the detailed functions and the molecular mechanisms of short Ago remain poorly understood.

Recent study has identified a prokaryotic immune system called short prokaryotic Argonaute TIR-APAZ (SPARTA) consisting of short Ago and TIR-APAZ in an operon, which protect their host from invading plasmid via TIR catalyzed NAD^+^ depletion ^[10]^. In the SPARTA system, short Ago originally binds TIR-APAZ forming a heterodimer, wherein the NADase activity of TIR-APAZ is repressed by short Ago. Upon the guide RNA-mediated target DNA binding, the SPARTA heterodimer assembles into higher-order oligomers where the TIR mediated NADase activity is unleashed ^[10]^. Unlike the well-characterized long Ago that typically functions as an endonuclease to silence the guide complementary target, short Ago in the SPARTA system likely serves as a nucleic acids sensor to control the NADase activity. However, how short Ago inhibits and activates accessory effector TIR in response to guide-target duplex binding remains unclear.

Toll/interleukin-1/ resistance gene (TIR) domain is broadly distributed across the tree of life as an essential component of immune systems ^[13–19]^. Generally, activation of TIR domains is oligomerization-dependent ^[19–25]^. Through self-assembly or association with other TIR-containing partners, eukaryotic TIR domain predominantly functions either as an enzyme hydrolyzing NAD^+^ or a scaffold facilitating signal transduction ^[26,27]^. Compared to the eukaryotic counterparts, the knowledge on structural assembly of prokaryotic TIR domains are limited. Recent study showed bacterial Sting and pyrimidine cyclase-associated TIR assembles into filament to complete NADase function in response to phage invading ^[20–23]^. In SPARTA system, it remains to be investigated how the TIR domain is arranged in the oligomerized SPARTA to fulfill the NADase activation.

Here, we focused on the working mechanism of SPARTA system from *Crenotalea thermophila*. Using cryogenic electronic microscopy (cryo-EM), we obtained a comprehensive set of structures representing the auto-inhibited state, three transient states as well as the active state of CrtSPARTA. Together with biochemical evidences, our results revealed detailed mechanism on auto inhibition and target ssDNA binding induced activation of SPARTA. A conserved C-terminal acidic tail (C-tail) of TIR-APAZ occupies the guide-target binding channel constituted by short Ago and APAZ to repress the SPARTA activity. This auto-inhibition would be unleashed upon expelling of the C-tail by the guide-meditated target binding. Appropriate guide-target duplex formation in channel triggers a series of conformational changes and thereafter induces the exposure of a new Ago-Ago dimerization interface to facilitate SPARTA oligomerization. The activated SPARTA revealed a TIR-APAZ_4_/short Ago_4_ octamer in which the four TIR domains are reorganized and packed to shape the NADase active center. Our findings form the structural basis for elaborated auto-inhibition and activation of SPARTA, and expand our understanding of short prokaryotic Ago in bacterial defense against invading nucleic acids.

## Result

### Reconstitution and cryo-EM structures of CrtSPARTA assemblies

To reconstitute the SPARTA system, we first purified a heterodimeric TIR-APAZ/short pAgo complex from *Crenotalea thermophila* (CrtSPARTA). CrtSPARTA incubated with 5′-phosphorylated (5′-P) 21 nt-long ssRNA guides and 25 nt-long complementary ssDNA targets showed little NADase activity at 37°C, whereas its activity was remarkably more efficient at 50°C **(Extended Data Fig.1a)**. We then reconstituted the SPARTA at 50°C and analyzed the oligomerization states using size exclusion chromatography (SEC), in accord with *Maribacter polysiphoniae* SPARTA ^[10]^, CrtSPARTA heterodimer also oligomerizes into octamer upon guide-target binding **(Extended Data Fig.1b)**.

To reveal the structural basis of SPARTA activation, we applied cryogenic electron microscopy (cryo-EM) to determine the CrtSPARTA structures in apo form and guide-target bound forms **(Extended Data Fig.2)**. In the guide-target bound dimer, we were able to classify two transient states, snapshotting the propagation of guide-target duplex **(Extended Data Fig.2-4)**. In the guide target bound octamer, we captured both octamer and tetramer forms, wherein TIR-APAZ and short Ago assemble in a stoichiometry of 4:4 and 2:2, respectively **(Extended Data Fig.2-4)**, with each heterodimer bound with guide-target duplex. In CrtSPARTA, TIR-APAZ consists of a N-terminal canonical TIR domain (1-138 aa), central APAZ domain (161-419 aa) and a C-terminal helical tail (C-tail, 420-450 aa). Short Ago is constituted with MID domain (1-268 aa) and a dead PIWI domain (269-507 aa), primarily responsible for guide RNA 5’ end recognition and nucleic acids binding respectively **(Fig.1a, Extended Data Fig.5a)**. Although named analog of PAZ, the fold of APAZ and its position relative to short Ago suggests it is actually equivalent to the N domain and the connected linker L1 of long Ago ^[28,29]^ **(Extended Data Fig.5b)**. The overall architecture of APAZ-short Ago resembles long Ago except for the absent PAZ domain **(Extended Data Fig.5c)** ^[30,31]^. APAZ and short Ago form two lobes to create a guide-target duplex binding channel, with the appended N terminal TIR domain extending from APAZ and reaching to MID domain of short Ago **(Fig.1b)**. The C-tail extends from APAZ and interacts with the TIR domain via the assembly of two β-strands. Notably, the C-tail is inserted into the guide-target binding channel **(Fig. 1b)**. The occupation of the guide-target binding channel by C-tail raises a hypothesis whether it regulates the SPARTA activity.

**Figure 1.**
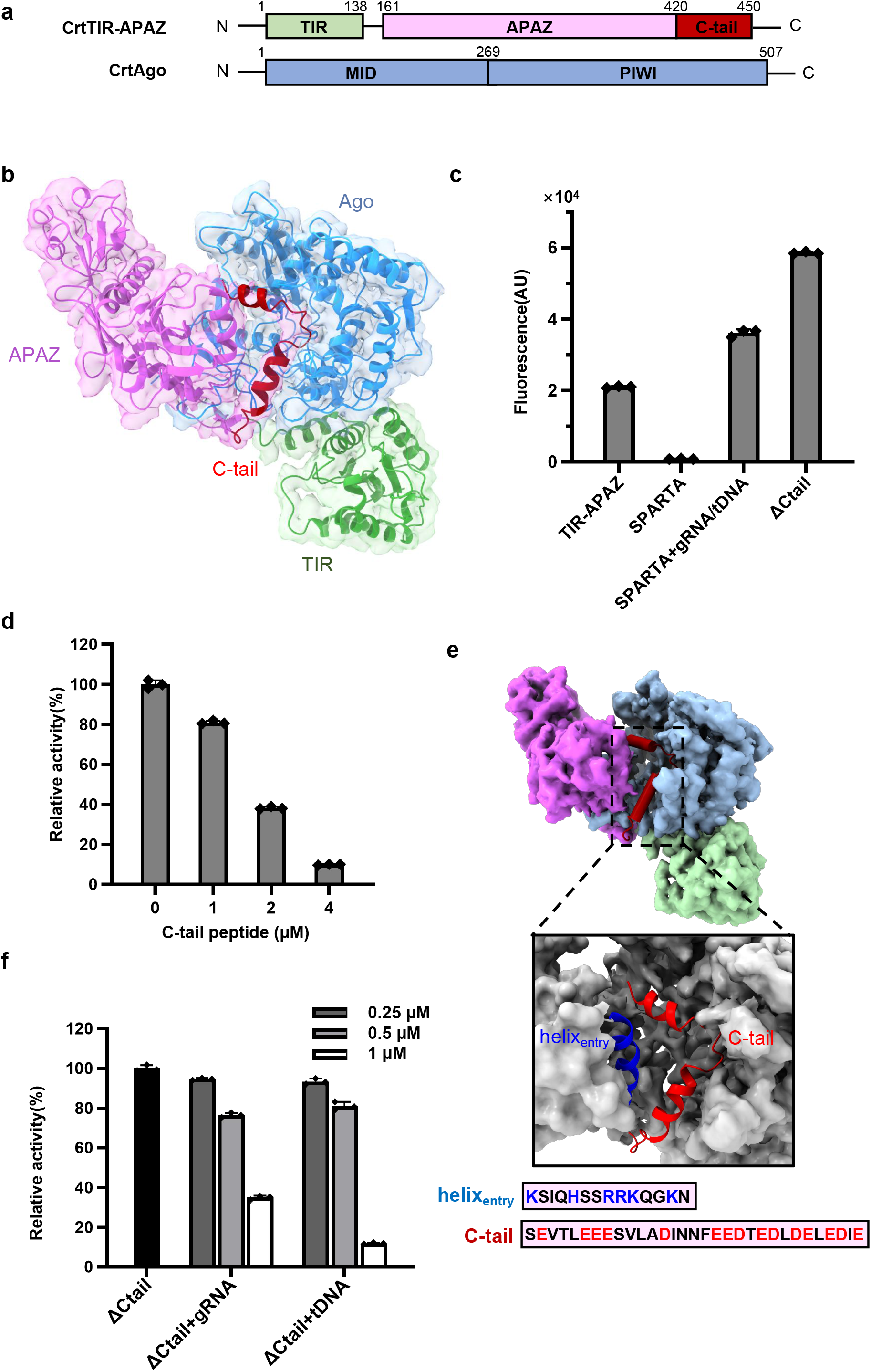
Auto-inhibition of SPARTA by an acidic tail. **a.** The primary structure of CrtTIR-APAZ and CrtAgo. **b.** The cryo-EM structure of CrtSPARTA apo form. The structure model is shown in cartoon with Cryo-EM map. The TIR, APAZ, and Ago are colored in green, purple, and blue respectively. The C-tail occupies the guide-target binding channel is indicated. **c.** The NADase activity of CrtSPARTA. The flurescene were measured for reactions catalyzed by CrtTIR-APAZ, CrtSPARTA with or w/o guide-target, and the C-tail truncated CrtSPARTA (ΔCtail), respectively. GT: guide RNA and target DNA. **d.** The inhibitory effect of supplemented synthetic C-tail in a dose dependent manner. The activity of the C-tail truncated SPARTA (ΔCtail) was normalized to 100%. **e.** The close-up view of the acidic C-tail extending along a basic helix to the channel. The amino acid sequences of the C-tail and the helix_entry_ are listed below with the acidic and basic residues highlighted in red and blue, respectively. **f.** The inhibitory effect of solely guide RNA or target DNA on C-tail truncated SPARTA. The activity of the C-tail truncated SPARTA (ΔCtail) was normalized to 100%.

### SPARTA is auto-inhibited by the acidic C-tail

To investigate the function of C-tail in SPARTA, we first checked the NADase activity of SPARTA in details. CrtTIR-APAZ alone is able to hydrolyze NAD^+^, however, this activity is quenched in the presence of short ago **(Fig.1c)**, which can be unleashed by guide-RNA-mediated target DNA binding. Surprisingly, the C-tail truncated SPARTA (SPARTA_ΔCtail) is able to efficiently catalyzes the NAD^+^ degradation **(Fig.1c)**. To confirm the resumed activity of SPARTA_ΔCtail is a direct result of the C-tail release from guide-target binding channel, we complemented the SPARTA_ΔCtail with synthetic C-tail peptide and checked the NADase activity. The results showed that addition of C-tail peptide would substantially reduce the NADase activity in a dose-dependent manner, and almost fully repress SPARTA_ΔCtail when the C-tail is 3-fold excess **(Fig.1d)**. Collectively, our data indicate that the C-tail functions as a strong repressor against innate NADase activity of SPARTA.

Notably, the entire C-tail is strongly acidic with dominant D/E residues **(Fig.1e)**, implying its competence to fit into the positively charged channel as a nucleic acids mimicry. Specifically, at the channel entrance, the C-tail extends along a basic helix of APAZ (helix_entry_: 353-367 aa) enriched in Lys, Arg and His **(Fig.1e)**, which may direct the acidic C-tail extension into the channel. Sequence alignment of 56 TIR-APAZ C-terminal regions, as well as the basic helix, suggests that the SPARTA systems may adopt a conserved mechanism to regulate their activity using an acidic C-tail **(Extended Data Fig.6)**. We therefore speculated that the plug-in C-tail may inhibit SPARTA activation in a manner that mimics and competes with guide-target, consistent with previous findings that SPARTA can only be activated in vivo by the multi-copy plasmids. Actually, we did observe that guide alone cannot stably bind to SPARTA wt, whereas the binding is greatly enhanced when the C-tail is removed, with the A260/280 value of complex raises from 0.8 to 1.2 **(Extended Data Fig.7)**. Furthermore, while SPARTA_ΔCtail with or without guide-target duplex is active, the addition of either guide or target alone to SPARTA_ΔCtail would abolish its NADase activity **(Fig. 1f)**, showing single strand nucleic acids possess similar repression effect as the C-tail to SPARTA. Together with the fact that wild type SPARTA would be activated by guided-mediated complementary target binding, these results suggest that either the fully evacuation or the “correct fit” of the guide-target binding channel is necessary for SPARTA activation.

### Guide-target duplex binding activates CrtSPARTA

The structures of SPARTA hetero<u>dimer</u> bound with <u>g</u>uide-target (Dimer-GT) provide detailed mechanism how the guide-target binding activates SPARTA. We captured two structural snapshots representing the transition states prior to activation, named Dimer-GT1 and Dimer-GT2, respectively **(Fig. 2a)**. Structural alignment of Dimer-GT1 and Dimer-GT2 with CrtSPARTA apo form shows the C-tail gradually moving away from guide-target binding channel: in Dimer-GT1, the C-tail is repulsed out from the guide-target binding channel but still hang on nearby, and is completely invisible in Dimer-GT2 **(Fig.2b)**. These observations confirm that C-tail is incompatible with guide-target and its releasing is necessary for SPARTA activation. The C-tail release is accompanied by the movement of helix_entry_, which withdrawals upon guide-target binding and contact the minor groove of guide-target duplex spanning nucleotide 5-10 **(Extended Data Fig.8a)**.

**Figure 2.**
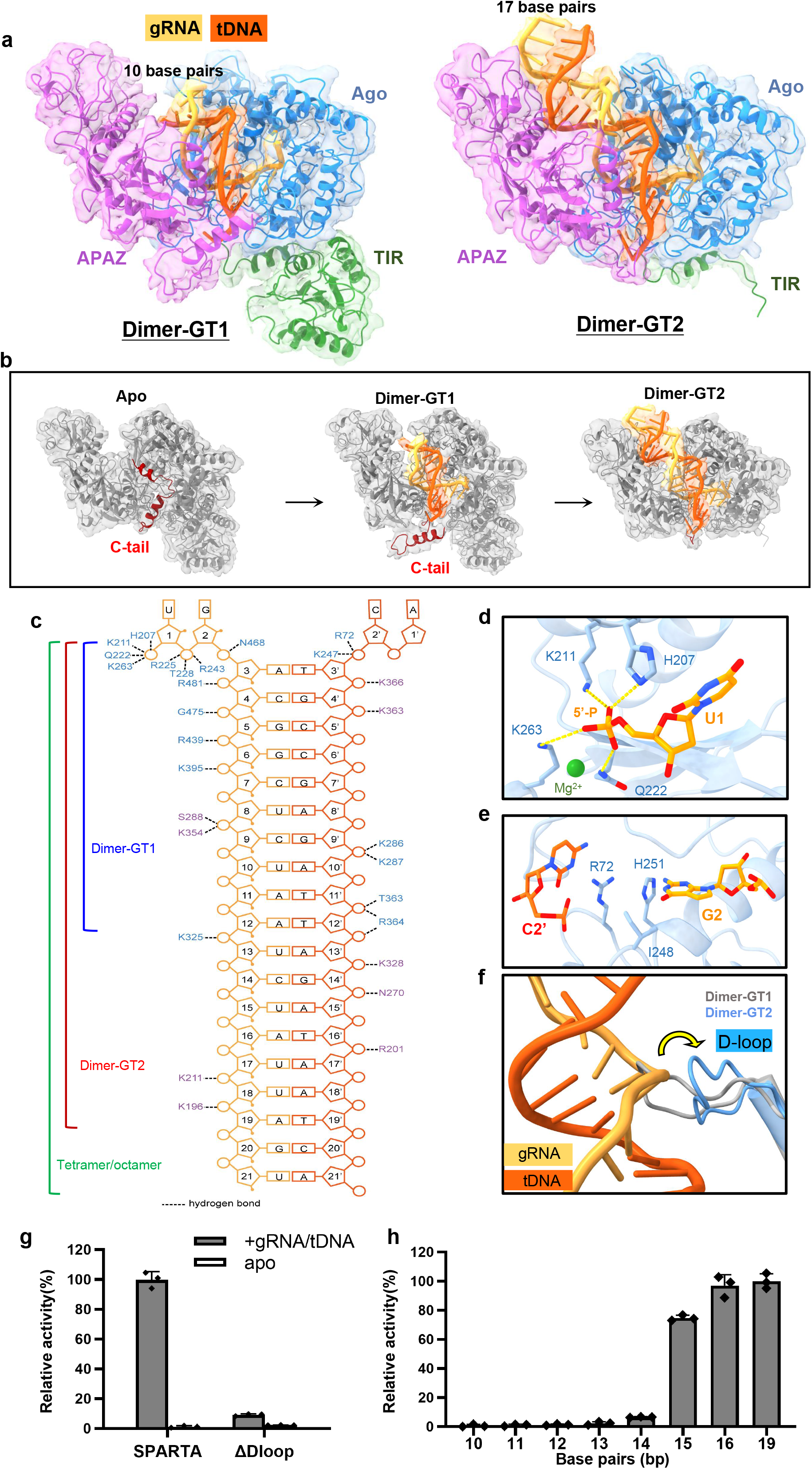
Guide-target duplex binding activates SPARTA. **a.** The structures of CrtSPARTA bound with guide RNA and target DNA in different states. *Left*, Dimer-GT1 with guide-target duplex of 10 base pairs; *right*, Dimer-GT2 with guide-target duplex of 17 base pairs. **b.** The conformational change of the C-tail along the binding and propagation of guide-target duplex. In Dimer-GT2, the C-tail is released and invisible in the structure. **c.** Schematic representation of the intermolecular contacts between SPARTA and guide-target in the complex structures. The base, backbone sugar, and phosphate group of the guide RNA (yellow) and target DNA (orange) are shown as rectangles, pentagons, and circles, respectively. Protein residues are color coded by domains. **d.** Recognition of the 5’P of guide RNA by the binding pocket in the MID domain of short Ago. **e.** The disruption of the base pair at the second nucleotide position. **f.** Superimposition of Dimer-GT1 (grey) and Dimer-GT2 (blue) showing the flipping of D-loop (residues 479–488, as indicated by the arrow) in the PIWI domain of short Ago upon guide-target duplex propagation from the 10 bp to the 17 bp duplex. **g.** The NADase activity of the D-loop truncated SPARTA (ΔDloop). The activity of the wild-type CrtSPARTA was normalized to 100%. **h.** Determination of the minimum length of the guide-target duplex required for CrtSPARTA activation. The flipped base 1 and 2 are not counted. The activity of the 19 bp-long guide-target duplex was normalized to 100%.

The Dimer-GT1 and Dimer-GT2 show overall similar architecture, as the guide-target duplex is embedded in the channel and the phosphate backbones extensively interacts with positively charged Arg and Lys spanning along the channel **(Fig. 2c)**. Akin to other prokaryotic Ago structures, in SPARTA the first uridine of guide flips and anchors in the MID pocket, with the 5’-P specifically recognized by the highly conserved residues H207, K211 Q222, and K263 **(Fig. 2d)**. Strikingly, unlike in other Ago structures that the guide and target starts base paired from the second nucleotides ^[5]^, this base pair is disrupted in SPARTA by R72, I248 and H251 of MID domain **(Fig. 2e)**, consistent with previous biochemical result that the mismatch at second nucleotide would not weaken and even slightly enhance the NADase activity of SPARTA ^[10]^.

Significantly, the structures of Dimer-GT1 and Dimer-GT2 reveal different guide-target binding status **(Fig. 2a)**. In Dimer-GT1, only density of 10 base pairs of guide-target duplex can be observed, spanning nucleotides 3-12 **(Fig.2a, Extended Data 8b)**. In contrast, guide-target duplex propagating to 17 bp can be assigned in Dimer-GT2, although the base pairs beyond 14 bp probably in transition are not perfectly aligned **(Fig.2a, Extended Data Fig.8b)**. Strikingly, we observed that a dynamic loop (D_loop: 318-330 aa of short Ago) gating the channel in SPARTA-apo undergoes conformational change upon guide-target duplex propagation **(Fig. 2f)**. Specifically, in Dimer-GT1, the D-loop closely contacts the nucleotide 12 of guide and blocks further guide-target base pair, whereas it flips out to allow for full guide-target base pairing in Dimer-GT2 **(Fig. 2f)**. Indeed, the conformational change of an equivalent loop was also observed in *Tt*Ago upon duplex propagation ^[32]^. We then asked if the D-loop movement is related to SPARTA activation. We superimposed the apo, Dimer-GT1, Dimer-GT2 and tetramer structures to snapshot D-loop movement, and along the transition it shows continuous and unidirectional shift of D-loop away from channel **(Extended Data Fig.8c)**, indicating that the activation of SPARTA is likely highly correlated with D-loop shift induced by guide-target binding.

To test this hypothesis, we evaluated NADase activity and the octamer formation of D-loop truncated SPARTA (SPARTA_ΔDloop). The results showed that the D-loop truncation disrupts the octamer formation and inactivates SPARTA **(Fig.2g, Extended Data Fig.8d)**, suggesting the essential role of the flipping D-loop probably as a trigger to activate the SPARTA through proofreading the guide-target complementarity. Then, we examined the minimum length of guide-target duplex required for D-loop extruding and thereafter SPARTA activation. The results showed that the wild-type SPARTA cannot be efficiently activated until the guide-target duplex extends to 15 bp long **(Fig.2h)**. We assume that the 15 bp long duplex may be stringently required for the complete repositioning of D-loop to the activation mode.

Besides the C-tail and D-loop, another obvious change driven by guide target binding is the releasing of TIR domain. In Dimer-GT1, just as the apo form, we were able to unambiguously assign each domain of SPARTA **(Fig. 2a left, Extended Data Fig.9a)**. However, in Dimer-GT2, the density of TIR domain is too weak to model, suggesting high flexibility of TIR domain upon full guide-target duplex formation **(Fig. 2a right, Extended Data Fig.9b)**. These evidences imply freeing TIR may be correlated with subsequent activation. Since N terminal TIR and C-tail closely contact with each other via two paralleled β-sheet, we speculated that the C tail repulse from guide-target binding channel may be companied by TIR release. We recognize that the shift of TIR from a fixed to a flexible state is crucial for the activation of SPARTA. This transformation allows TIR to reorient for oligomerization, which is also observed in other TIR-containing systems ^[24,25,33–35]^ (See below: SPARTA tetramer and octamer).

Collectively, our two guide-target bound structures suggested that nucleic acids binding excludes C-tail, releases TIR, and repulses the D-loop. These conformational changes possibly trigger the SPARTA activation.

### Ago-Ago interaction oligomerizes CrtSPARTA

Following complete guide-target duplex binding, the two heterodimers form a “Butterfly” shaped tetramer that is mediated by a novel Ago-Ago dimerization interface **(Fig.3a)**. Extensive interactions were observed at the interface resulting in a buried area of 1272 Å^2^. This inter-strand interaction primarily involves three regions of Ago residues, including residues 35-40, 129-137, and 498-504 **(Fig.3a)**. The residues Y37 and K40 form hydrogen bond with K85’ and Q35’, respectively. At another side, highly intensive electrostatic interactions are observed between the consecutive polar residues 129-NKNDEE-134 and residues N34’/Y262’/K267’/T498’/L501’/K504’/Y505’, while R135 and D137 crossly interact with each other from the adjacent protomer **(Fig.3a, right)**. The interaction is doubly reinforced through two-fold symmetry of Ago-Ago. To verify the structure and the importance of the interactions for SPARTA activation, we introduced mutations at these residues. The mutation of E133A/R135A/D137A (ERD^AAA^) reduced octamer formation, and substantially impaired NADase activity. While the mutation of Y37A/K40A (YK^AA^) entirely abolished NADase activity and complete disrupted octamer **(Fig.3b, Extended Data Fig.10)**. These results show the correlation between SPARTA activation and octamer formation. By contrast, both the mutants retain the guide-target binding ability comparable to wild type **(Extended Data Fig.10)**, suggesting the inactivation is the result of oligomer disruption at the Ago-Ago interface, but not the other factors caused by nucleic acids binding. The residual activity and slight oligomer formation for E133A/R135A/D137A could be explained by the strong interaction cluster involving nearly twenty pairs interaction in this region. Therefore, the SPARTA oligomerization is mediated by the strong Ago-Ago interface, which is essential for SPARTA activation.

**Figure 3.**
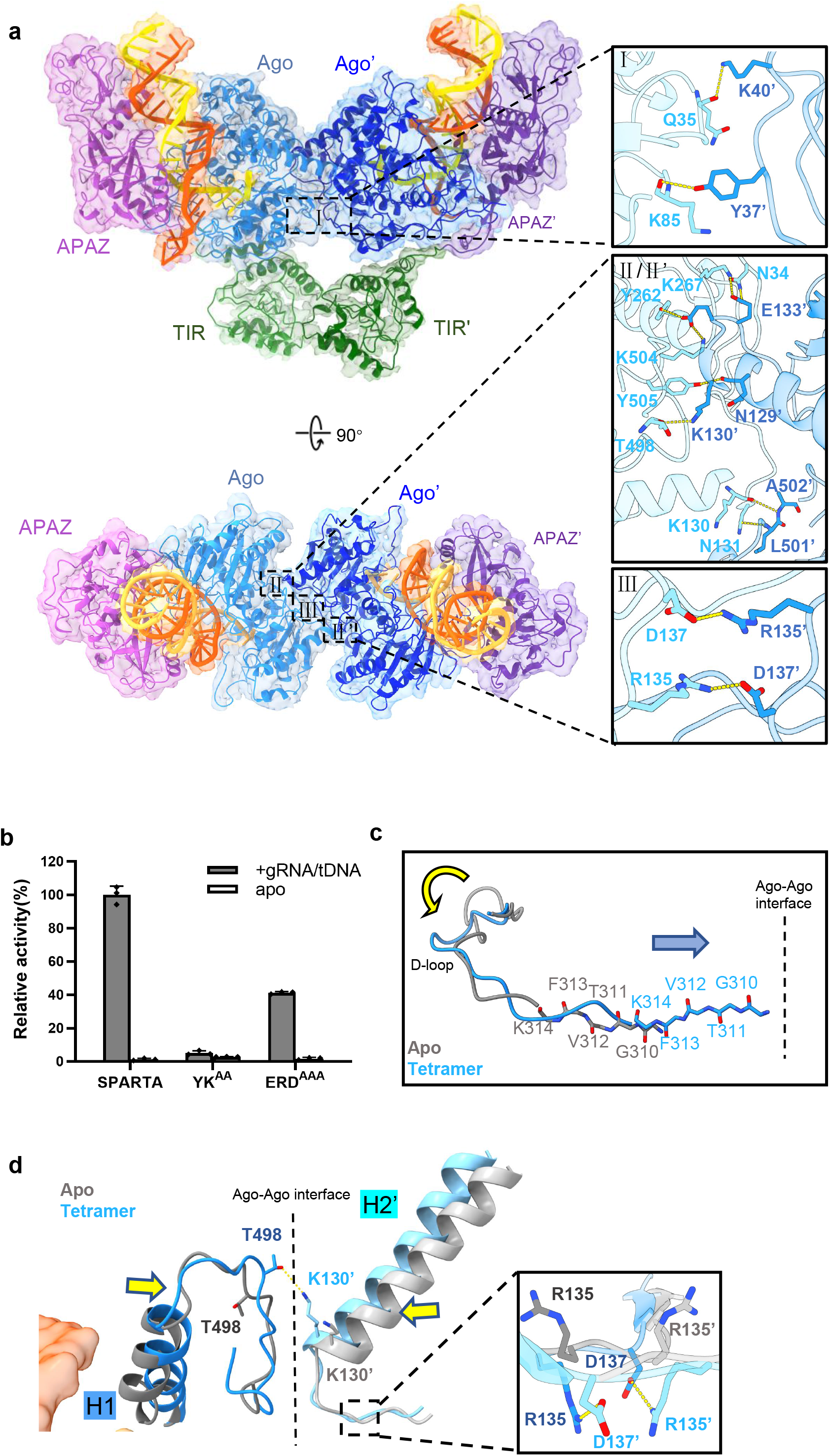
Ago-Ago interaction induced by guide-target binding is the prerequisite for CrtSPARTA activation. **a.** Two orthogonal views of CrtSPARTA tetramer showing the extensive interactions at the Ago-Ago dimerization interface. The insertions show the three clusters of interactions. **b.** The NADase activity of the Ago-Ago interface mutants. YK^AA^: Y37A/K40A; ERD^AAA^: E133A/R135A/D137A. The activity of the wild-type CrtSPARTA was normalized to 100%. **c.** The three-residue sliding of the D-loop connected β-strand (G310 to K314 of short Ago) towards the Ago-Ago interface direction upon guide-target duplex binding. **d.** Superimposition of CrtSPARTA apo form (Ago_a_ in dark grey, Ago_b_ in light grey) and tetramer (dark blue, light blue) showing the substantial conformational changes at Ago-Ago interface. The transition from the apo form to the tetramer state involves the movement of helices H1 and H2, enabling hydrogen bond formation between T498 and K130’ at the interface in tetramer. The insertion shows the reposition of the residues R135 to interact with D137 from the adjacent Ago chain. The Ago-Ago interface is delineated by the dashed line.

To reveal the detailed mechanism that couples the guide-mediated target recognition and the Ago-Ago dimerization, we superimposed Dimer-apo onto tetramer structure to visualize the conformational change of Ago upon guide target binding. Besides D-loop, multiple regions adjacent to guide-target binding and Ago-Ago interface have obvious movements. The flipped D-loop forces the connected β-strand (310-314 aa of short Ago) to slide by three residues towards Ago-Ago interface direction **(Fig.3c)**, akin to one-residue sliding of the equivalent β-strand observed in long Ago structure ^[32]^. Noticeably, beneath the D loop and β-sheet, the helix H1 (478-492 aa) close to nucleotide 9 and 10 of target DNA shifts towards Ago-Ago interface **(Fig.3d)**. Such movement brings the attached loop (493-506 aa) in close proximity to the long helix H2 (T110-K130 aa of short Ago) of the adjacent Ago protomer, together with the forward movement of the helix H2 for approximate 2.3 Å in average, enables inter-chain hydrogen bond formation between T498 and K130’ at the Ago-Ago interface **(Fig.3d)**. The movement of the helix H2 further drags the connected loop (N131-D137 of short Ago) that greatly contribute to Ago-Ago dimerization via intensive interactions, where R135 is repositioned in tetramer to make cross contact with D137’ via near 2-fold symmetry **(Fig.3d)**. The shift of this essential loop, as well as the nearby regions probably initiates the shaping of Ago-Ago dimerization interface. Taken together, the Ago-Ago dimerization interface exposure is a result of synergetic conformational changes of short Ago in cascade, triggered by the guide-target binding.

### TIR rearrangement and packing shapes NAD^+^ active sites in SPARTA octamer

Examination of tetramer and octamer structures explains why TIR releasing and Ago-Ago dimerization interface formation are essential for SPARTA activation. In the SPARTA tetramer, the two TIR domains are near paralleled aligned with one another in a head-to-tail fashion **(Fig.4a)**. As a result, the DD loop (105-122 aa) from TIR_a_ and the BB loop (35-46 aa) from TIR_b_, stack on top of one another and thus shape one active site for NAD^+^ **(Fig.4a, Extended Data Fig.11a)**. To achieve the stacking interaction, one TIR from Dimer-apo has to turn 180° over via the flexible linker (139-159 aa) to align with the other TIR in the tetramer **(Fig. 4b)**, which makes our observation rational that freeing TIR in Dimer-GT2 is a prerequisite for tetramer formation. Subsequently, the Ago-Ago interaction brings the free to aligned TIRs into proximity to shape the active site formed by BB loop and DD loop.

**Figure 4.**
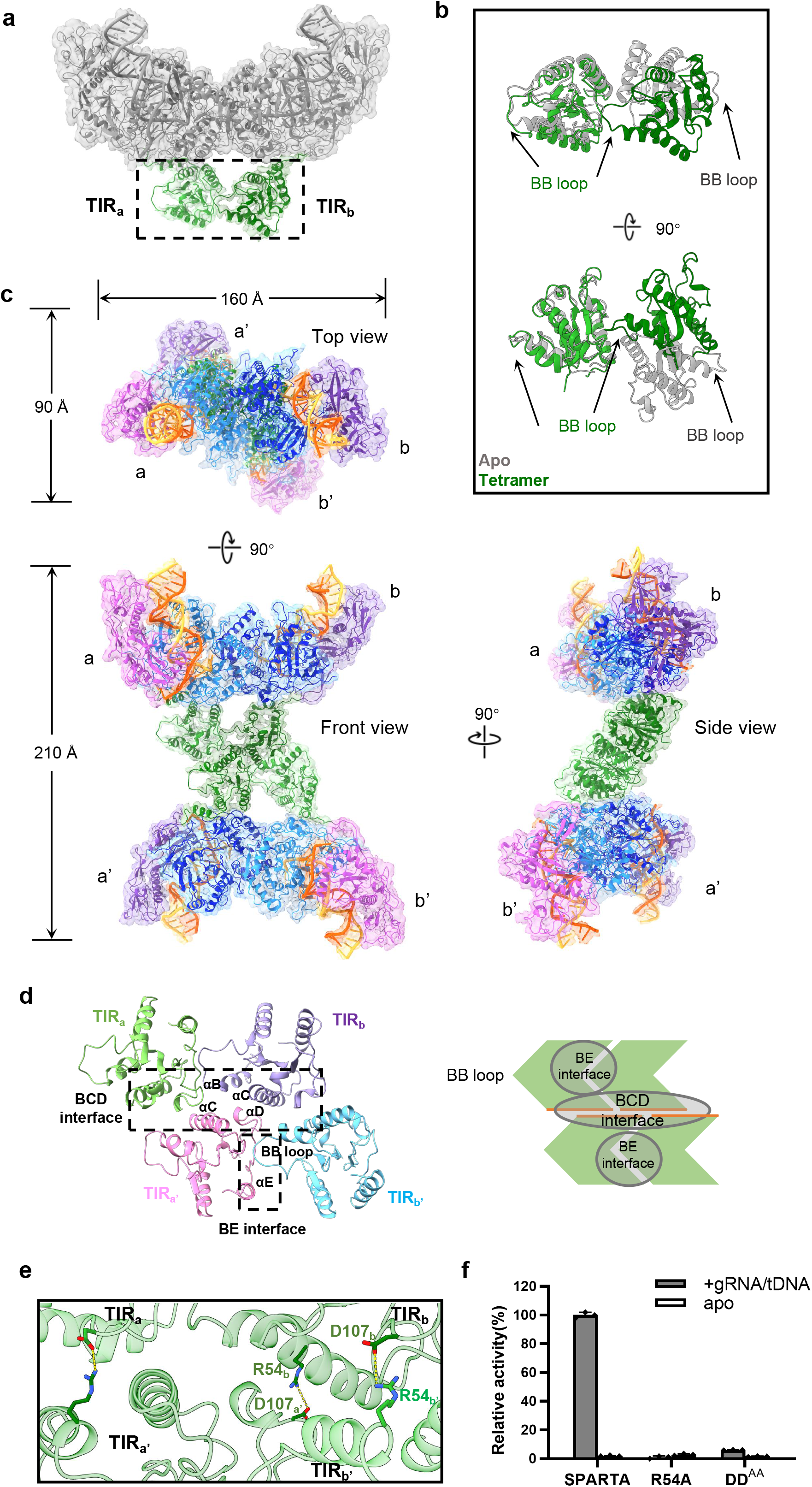
TIR rearrangement and packing shapes NAD^+^ active sites in SPARTA octamer. **a.** The head-to-tail arrangement of two TIR domains in the SPARTA tetramer. **b.** Superimposition of SPARTA apo (grey) onto tetramer (green) showing significant rearrangement of one TIR domain with an approximately 180° rotation. Two orthogonal views of the superimposed TIR pair are shown. **c.** Three orthogonal views of CrtSPARTA octamer structure. In octamer, each heterodimer bound with guide-target is labeled as a, a’, b, b’, respectively. **d.** Four TIR domains arranged in the “scaffold assembly” via BE and BCD interface. The structure components constitute the interfaces are labeled. For clarity, the four TIR domains are shown in different colors, and a cartoon diagram was used to highlight the parallel arrangement and packing of TIR domains via BE and BCD interface. **e.** The R54-D107 interaction pairs at BCD interface of TIR domains. R54 interacts with D107 at the interface of TIR_a_-TIR_a’_, TIR_b_-TIR_b’_, and TIR_b’_-TIR_a_. **f.** The NADase activity of BCD interface mutants. DD^AA^: D106A/D107A. The activity of the wild-type CrtSPARTA was normalized to 100%.

Two tetramers further associate to form an octamer, with total dimensions of 160 Å X 90 Å X 210 Å **(Fig.4c)**. Two sets of head-to-tail TIR dimer are crossly paralleled packed at the center of SPARTA octamer, with two APAZ-Ago heterodimers tilted to each other **(Fig.4c, top view)**. The assembly of central four TIR domains, called “scaffold assembly”, involves two types of asymmetric interactions **(Fig.4d)**, and similar packing was also observed in MAL-TIR ^[36]^ **(Extended Data Fig.11b)**, an adaptor in Toll-like receptor signaling. One is mediated by BE interface initiated in SPARTA tetramer, including the area around BB loop of TIR_a_, and DD loop, αE, βD and βE of TIR_b_. Another asymmetric interaction at BCD interface involves residues in the αB and αC helices of TIR_a_ (TIR_b_) and the αD helix and the CD loop of TIR_a’_ (TIR_b’_), scaffolding SPARTA tetramer into octamer **(Fig.4d, Extended Data Fig.11a)**). Unlike MAL-TIR that mainly relies on hydrophobic interactions at the BCD interface, this interface is enriched in D/E and K/R residues in SPARTA, including R54, E72, K83, K85, K86, D106, and D107, which possibly contributes to octamer formation via electrostatic interaction at the BCD interface. Particularly, at the BCD interface, three pairs of R54 and D107 were observed to interact with each other in TIR_a_-TIR_a’_, TIR_b_-TIR_b’_, and TIR_b’_-TIR_a_, respectively **(Fig.4e, left)**. To determine whether these interactions contribute to the formation of active SPARTA octamer, we mutated R54 and D106/D107 into Ala, respectively. The result shows that both the mutations of R54A and D106A/D107A (DD^AA^) would dramatically reduce the NADase activity of SPARTA **(Fig.4f)**, confirming the importance of these residues for active SPARTA formation. Because R54 is far away from NAD^+^ catalytic pocket, the abolished NADase activity of the mutant R54A should be a result of the disrupted BCD interface, which is unable to form functional octamer to stabilize the NAD^+^ active sites. Notably, the TIR density in octamer is better defined than in tetramer (**Extended Data Fig.9c,d)**, which also implies that TIRs are more stable in octamer via interactions at BCD interface. This evidence suggests that the “scaffold assembly” of four TIR domains is essential for the NADase activity of SPARTA.

Taken together, TIR domain rearrangement and their close packing result in the SPARTA octamer formation, shaping and stabilizing two dimer-mate NAD^+^ active sites.

## Discussion

Based on structural and biochemical results, we propose the mechanisms of SPARTA auto-inhibition and activation **(Fig.5)**: SPARTA is originally inactive due to the auto-inhibition by a conserved acidic C-tail blocking the guide-target binding channel, which can be released by the binding of guide-target duplex. Besides the release of inhibition, the binding and propagation guide-target duplex would trigger a series of conformational changes of SPARTA, including the D-loop flipping, Ago-Ago dimerization interface shaping, TIR rearrangement. These conformational changes ultimately result in the formation of activated higher-order assembly of SPARTA, wherein four TIR are reorganized to shape the NADase active sites.

**Figure 5.**
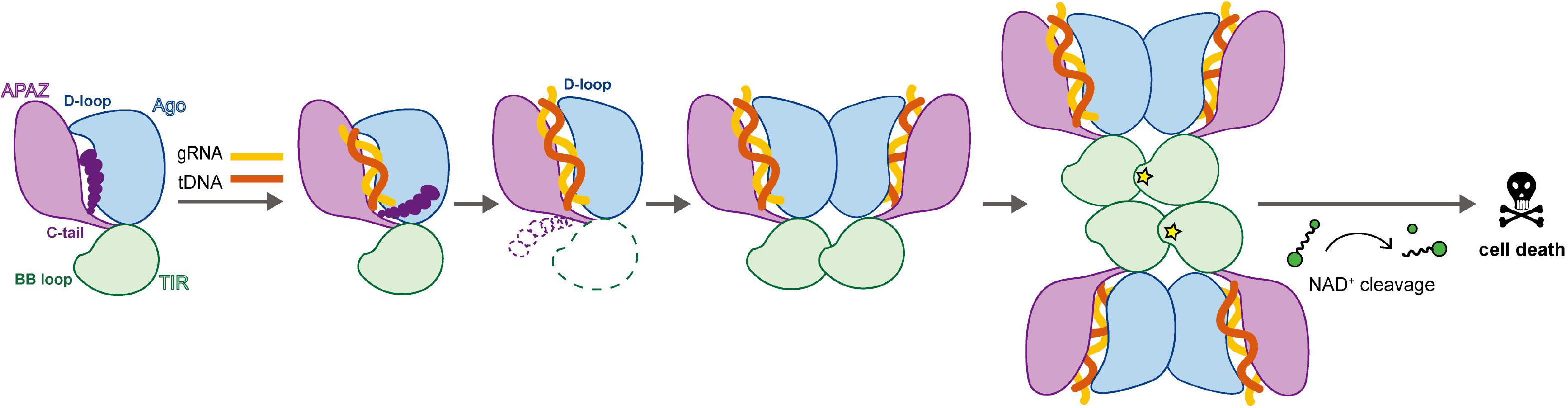
The proposal for the mechanism of SPARTA auto-inhibition and activation. SPARTA is auto-inhibited by an acidic C-tail, which blocks the guide-target binding channel constituted by the APAZ/short Ago. The guide-mediated sensing of target DNA releases the inhibition by repelling the C-tail and triggers a series of conformational changes of SPARTA, such as D-loop flipping and TIR freeing. These conformational changes induce Ago-Ago dimerization and ultimately result in the formation of activated SPARTA octamer, wherein four TIR are reorganized to shape the NADase active sites. The activated SPARTA depletes cellular NAD+, thereby mediating population-based immunity via the abortive infection mechanism.

Although the overall architecture resembles long Ago, the APAZ/short Ago as a regulator in SPARTA defense system differs significantly regarding the utilization of repressor, the guide-target binding, as well as the Ago oligomerization state.

The acidic C-terminal tail is conserved in SPARTA but absent in long Ago as well as short prokaryotic Argonaute Sir2-APAZ (SPARSA), suggesting its unique role in SPARTA auto-inhibition. Besides, the acidic mimicry of nucleic acids implies strong competition between C-tail and guide-target for the positively charged binding channel, which is supported by our binding assay as well as the previous finding that SPARTA can only be activated by a multi-copy plasmid, not by a low-copy one ^[10]^. This “design” enables SPARTA activation under stringent control to avoid either mis-activation by the partial complementary nucleic acids or over-activation by low-copy plasmid invading. We hypothesize that the formation of active SPARTA requires all four heterodimers bound with guide-target duplex, and this may be another strategy to minimize the mis-activation and over-activation.

Followed by inhibition unleashing, the guide-mediated target recognition triggers SPARTA activation. Our results showed that the flipping of D-loop upon the guide-target duplex propagation is essential for SPARTA activation, which is likely to function as a gating mechanism to proofread guide-target complementary. Earlier study has also observed the conformational change of an equivalent loop L1 and an accompanied loop L2 in *Tt*Ago induced by propagation of guide-target duplex, which couples the release of guide 3’ end from PAZ with the subsequent activation of PIWI for target cleavage ^[32]^.

Interestingly, the movement of loop L2 forces a glutamic acid residue inserting into the catalytic pocket to initiate PIWI cleavage activity ^[37]^. Such conformational change of the equivalent loop was unnecessary for catalytic inactive short Ago and was not observed in SPARTA. The lack of PAZ and the presence of “dead” PIWI suggests D-loop may functions differently in SPARTA. Actually, the D-loop is longer than loop L1, and it appears to have more profound effect on adjacent regions as extensive conformational changes were observed, which are probably necessary for Ago-Ago dimerization interface shaping and TIR’s NADase activation in a distance. These together implies that APAZ/short Ago likely repurpose the canonical structural elements of long Ago for SPARTA activation.

Besides the canonical elements of Ago, a new Ago-Ago dimerization interface is also critical for SPARTA function. Long Ago generally exists as a monomer. The crystal structures of the “truncated” Ago from *Archaeoglobus fulgidus* (*Af*Ago) consisting of MID and PIWI domain show close contacts between two Ago molecules (PDB ID: 1YTU, 6XUP) ^[38,39]^, and recent study confirms homodimer formation of *Af*Ago in solution ^[40]^, although the functional relevance of the dimer form is unknown. In this study, we observed a new Ago-Ago dimerization interface that serves as a scaffold to facilitate SPARTA oligomerization and activation. Whether the function-unknown *Af*Ago and other Ago-mediated defense systems also adopt similar mechanism to regulate activity requires further investigation ^[11,12,41,42]^.

TIR activation is generally driven by oligomerization, as the associated domains oligomerize upon sensing stimulus and further bring the TIR domains into proximity to assemble with each other ^[18–24]^. Correlated with the different functional roles, eukaryotic TIR domains are primarily arranged either as “scaffold assembly” or “enzymatic assembly”, corresponding to parallel or anti parallel two-stranded assembly via distinct interfaces, respectively (**Extended Data Fig.11b,c)** ^[24–25]^. So far, structurally characterized TIR-containing proteins show that “enzymatic assembly” formation of TIRs with NADase function (eg. TIR-NLR ^[23,29]^, SARM1 ^[30,31]^, TIR-Sting ^[19]^) is driven by homolateral oligomerization of the domains connected to TIR. By contrast, in SPARTA, guide-target binding stimulates Ago-Ago dimerization which brings two TIRs into proximity to form dimer first. Then, two TIR dimers are likely spontaneously stack into a tetramer. This spontaneous assembling results in an unusual “scaffold assembly” of TIR domains, other than “enzymatic assembly”, although it functions as a NADase. Given the diversity of prokaryotic TIR domains especially contextualized with its diverse partners, we may anticipate other new assembly forms of TIR correlated with different functions.

## Methods

### Protein expression and purification

The genes encoding *Crenotalea thermophila* TIR-APAZ and short Ago were cloned into the vector pCOLADuet-1 using homologous recombination, with the 6*His tag attached to the N-terminus of TIR-APAZ. For the TIR-APAZ alone, the gene was cloned into vector pET-28a. The constructs were transformed into the *E. coli* strain BL21 star (DE3) and the cell cultures were grown at 37°C to OD_600_ of 0.6. The protein expression was induced with addition of 0.25 mM isopropyl-β-D-thiogalactoside (IPTG) at 18°C for 20 h. The cells pellets were harvested and lysed by sonication in buffer containing 50 mM HEPES pH 8.0, 500 mM NaCl, and 5 mM imidazole. The supernatant was incubated with Ni-NTA resin pre-equilibrated with lysis buffer. The unbound proteins were washed out using lysis buffer containing 5, 20, and 50 mM imidazole respectively. The target proteins were eluted in the buffer containing 50 mM HEPES pH 8.0, 500 mM NaCl, and 300 mM imidazole. The eluted proteins were then loaded onto a Heparin column to remove nucleic acids and impurities. The TIR-APAZ/short Ago complex was further purified by gel filtration chromatography using a HiLoad 16/600 Superdex 200 prep grade column (Cytiva) equilibrated with 20 mM HEPES, pH 7.5, 500 mM NaCl. The purity was analyzed by SDS-PAGE and the fractions of interest were combined. The protein was concentrated, aliquoted, and stored at -80°C until use. The CrtSPARTA mutants followed the same procedure of expression and purification.

### Site-directed mutagenesis

All the mutations or the truncations of CrtSPARTA were introduced using ClonExpress II (Vazyme) with pCOLADuet-1_CrtSPARTA as template. All constructs were verified by DNA sequencing.

### In vitro NADase assay

The purified 1 μM CrtTIR-APAZ or CrtSPARTA was mixed with 1 μM 21 nt-long 5’-phosphorylated guide RNA and 25 nt-long target ssDNA. Reactions were then initiated by adding ε-NAD^+^ to a final concentration of 25 μM and incubated at 50°C for 30 min. After that, fluorescence emission at 410 nm of the reactions was read using a SPARK Multi-Mode Reader (TECAN) after excitation at 310 nm. Plots were generated with GraphPad Prism 9.3.0. To determine the C-tail’s function, the CrtSPART_ΔC-tail in the absence of guide RNA and target DNA was incubated with 25 μM ε-NAD^+^. To investigate the inhibitory effect of C-tail, RNA guide and DNA target on C-tail-truncated SPARTA (CrtSPARTA_ΔCtail), each of them was separately pre-incubated with CrtSPARTA_ΔCtail at 50°C for 10 min, after which the 25 μM ε-NAD^+^ was added and incubated at 50°C for 30 min. To determine the minimum length of guide-target duplex for SPARTA activation, target DNA in different length was incubated with SPARTA and 25 nt-long guide RNA, and followed the same procedure as above. The experiments were repeated in triplicates, and the measurements were taken from distinct samples. The data are expressed as the mean ± standard deviation (SD). The sequence of guide RNA and target DNA used in activity assay were summarized in Extended Data Table 1.

### CrtSPARTA assembly analysis by size exclusion chromatography

10 μM purified CrtSPARTA or its mutants was incubated for 5 min at 50°C with guide RNA in 1:1.2 molar ratio in buffer containing 20 mM HEPES pH7.5, 150 mM KCl, 5 mM MnCl_2_, 10% glycerol. Thereafter, target DNA was added in a 1:1.2 molar ratio and samples were incubated at 50°C for 30 min. After incubation samples were loaded onto a Superdex 200 Increase 10/300 GL column (Cytiva Life Sciences) and different oligomer states were resolved. The absorption at 280nm and 260nm were monitored.

For preparation of Cryo-EM sample, the peaks of guide-target bounded dimer and octamer were collected separately, and concentrated to a proper concentration for Cryo-EM experiments.

### Electron microscopy sample preparation and data acquisition

3.5 μL of purified *Crt* SPARTA complex at a concentration of 0.5 mg/mL was applied to glow-discharged Quantifoil holey gold grids (R 1.2/1.3, Au 300 mesh) followed by a 15 s wait time and 5 s blot time with the blot force set to 2. Then, the grids plunge-frozen in liquid ethane using a Virtrobot Mark IV (Thermo Fisher). The Vitrobot chamber was maintained at close to 100% humidity and 22°C. Cryo-EM images were manually collected on a FEI Titan Krios G3i (Thermo Fisher) operated at 300 kV and equipped with a K2 Summit direct electron detector. All cryo-EM movies were recorded in counting mode with SerialEM. The detailed parameters including electron dose, pixel size and magnifications of electron microscopy data collection are listed in Extended Data Table 2.

### EM data processing

Images were processed in cryoSPARC (v.4.2) unless indicated otherwise. Movie frames were aligned and summed using Motioncor2. Contrast transfer (CTF) parameters were estimated for individual particles on each micrograph using cryoSPARC. After patch CTF estimation, micrographs with a resolution estimation worse than 6 Å were discarded. Blob picker was used to pick particles, using a circular diameter of 60 to 240 Å. Particle picks were inspected and particles with NCC scores below 0.3 were discarded. 2D classification, 3D classification and 3D refinement were also carried out using cryoSPARC. All refinements followed the gold-standard procedure, in which two half data sets were refined independently. The overall resolutions were estimated based on the gold-standard criterion of Fourier shell correlation (FSC) = 0.143. Local resolutions were calculated in cryoSPARC and visualized using ChimeraX.

For the [Data set 1, Apo-form], 3408 movies were recorded, and 3395 images were kept after manual inspection. After 2D classification, 1,641,247 particles out of 3,713,753 auto-picking particles were retained and subjected for 3D reconstruction and 3D classification. To improve the resolution, 639,290 particles belonging to the best two 3D classes were retained for another round of 3D refinement, and 375,800 particles in one best class were retained. Further 3D refinement, followed by non-uniform refinement, local refinement and post processing, resulted in a 3.3 Å average resolution 3D map.

For the [Data set 2, Dimer-GT1 and Dimer-GT2], 3199 movies were recorded, and 3182 images were kept after manual inspection. After 2D classification, 1,156,841 particles out of 4,327,470 auto-picking particles were retained and subjected for 3D reconstruction and 3D classification. To improve the resolution, 405,852 particles belonging to the best two 3D classes were retained for another round of 3D refinement, and 156,464 particles in one best class were retained. Further 3D refinement, followed by non-uniform refinement, local refinement and post-processing, resulted in a 3.5 Å average resolution 3D map [Data set 2, Dimer-GT1]. In addition, 117,819 particles belonging to a class with incomplete density and no apparent orientation preference from the initial 3D classification were selected for further 3D refinement and post-processing, resulted in a 3.4 Å average resolution 3D map [Data set 2, Dimer-GT2].

For the [Data set 3, SPARTA tetramer and octamer], 3626 movies were recorded, and 3620 images were kept after manual inspection. The 2D classification results revealed two different sizes of particles. Among the 7,596,546 automatically picked particles, a subset of 1,069,070 particles identified as being in the tetramer state, as well as an additional subset of 436,675 particles in the octamer state, were individually selected for subsequent 3D reconstruction and 3D classification. After multiple rounds of 3D classification. Finally, a 3.4 Å average resolution 3D map of [SPARTA tetramer] was generated using 174,896 particles through 3D refinement, non-uniform refinement, local refinement and post-processing. Similarly, a 4.2 Å average resolution 3D map of [SPARTA octamer] was generated using 147,489 particles through 3D refinement, non-uniform refinement, local refinement and post processing.

The detailed data processing and refinement statistics for the cryo-EM structures are summarized in Extended Data Table 2.

### Model building

The structures of the Crtshort Ago and CrtTIR-APAZ proteins (accessions in GenBank: WP_092459742.1, WP_092459739.1) were individually predicted using AlphaFold2, and the best ranking AlphaFold2 predicted structures out of five predictions were chosen as models.

The model was rigid-body fitted into the density using UCSF ChimeraX and manual rebuilding in Coot. Nucleotides and amino acid residues were mutated to reflect the true sequence of the construct. Models were subsequently refined with phenix.real_space_refine, using reference to the starting structures and restraints on protein secondary structure, nucleobase pairing, and nucleobase stacking. The quality of the structural model was checked using the MolProbity program in Phenix.

### Data availability

Atomic coordinates, maps and structure factors of the reported cryo-EM structures have been deposited in the Protein Data Bank under accession numbers 8J84 (Short ago complexed with TIR-APAZ), 8J9G (CrtSPARTA hetero-dimer bound with guide-target, state 1), 8J8H (CrtSPARTA hetero-dimer bound with guide-target, state 2), 8J9P (CrtSPARTA tetramer bound with guide-target) and 8JAY (CrtSPARTA octamer bound with guide-target) and in the Electron Microscopy Data Bank under accession codes EMD-36059 (Short ago complexed with TIR-APAZ), 36095 (CrtSPARTA hetero-dimer bound with guide-target, state 1), 36070 (CrtSPARTA hetero-dimer bound with guide target, state 2), 36114 (CrtSPARTA tetramer bound with guide-target) and 36138 (CrtSPARTA octamer bound with guide-target). Source data are provided with this paper.

## Acknowledgements

We would like to thank the Instrument Analysis Center (IAC) at Shanghai Jiao Tong University for Cryo-EM data collection. This work is supported by National Key Research and Development Program of China (2018YFA0902000), STI2030-Major Projects (2021ZD0203400), National Natural Science Foundation of China (31970547, 32271330, 32000889), Natural Science Foundation of Jiangsu Province (No.BK20190552), the Project Program of State Key Laboratory of Natural Medicines, China Pharmaceutical University (No.SKLNMZZ202014), and the Fundamental Research Funds for the Central Universities (No. 2632023GR17).

## Author contributions

M.R.C. and Y.B.X. conceived the project and designed the experiments. L.J.G., P.P.H., Z.X.L., and P.R.Y. carried out the experiments. M.R.C., Y.B.X., L.J.G., P.P.H., Z.X.L., Y.C.S., and M.L.L. analyzed the data. M.R.C, Y.B.X, P.P.H, and Z.X.L. wrote the manuscript. All authors discussed the results and contributed to the final manuscript.

## Figure legend

**Extended Data Fig 1.**
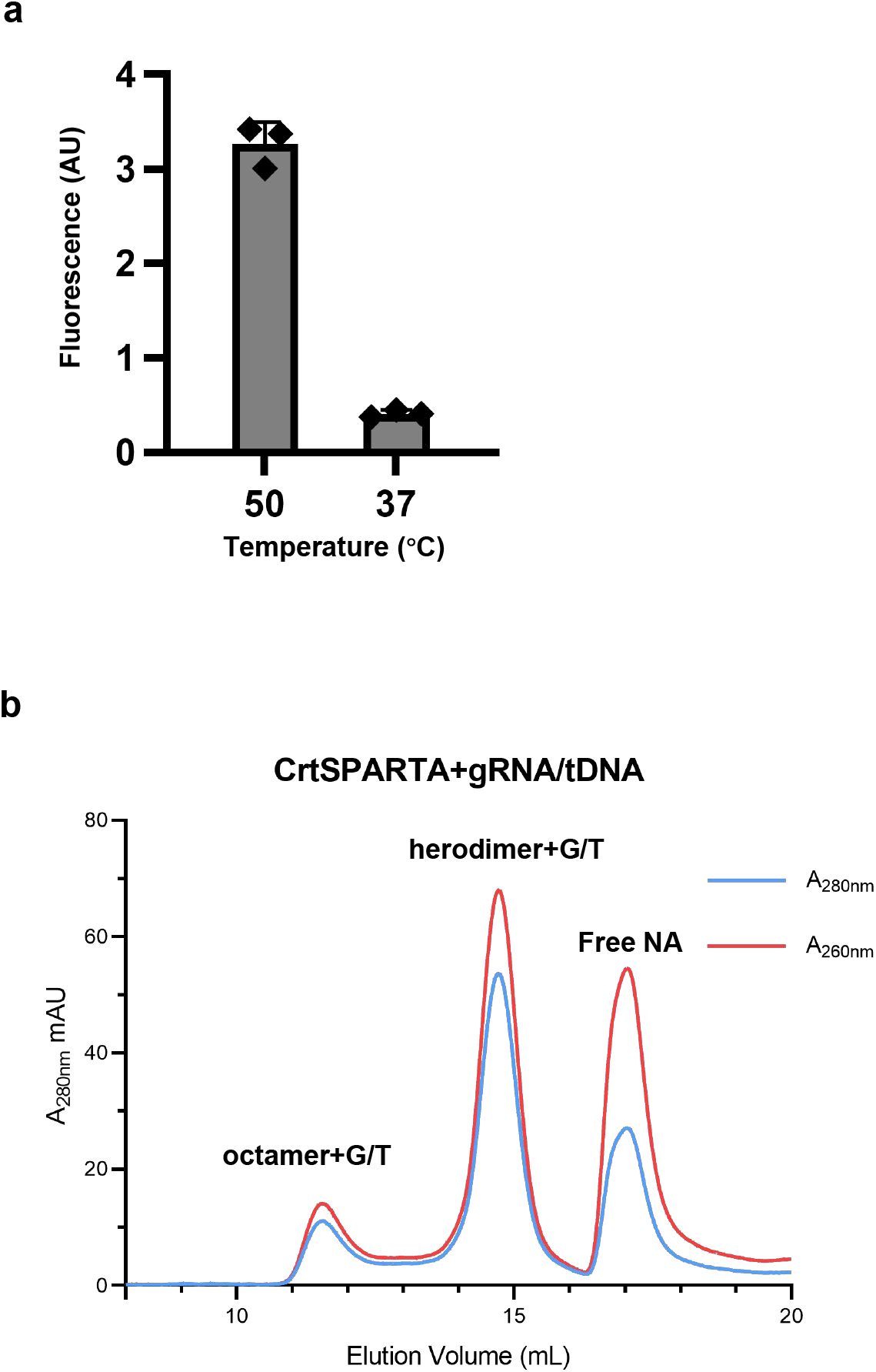
The reconstitution of CrtSPARTA complexes. a.NADase activity at 37 ° C and 50 ° C. b. The size exclusion chromatography of reconstituted CrtSPARTA complexes.

**Extended Data Fig 2.**
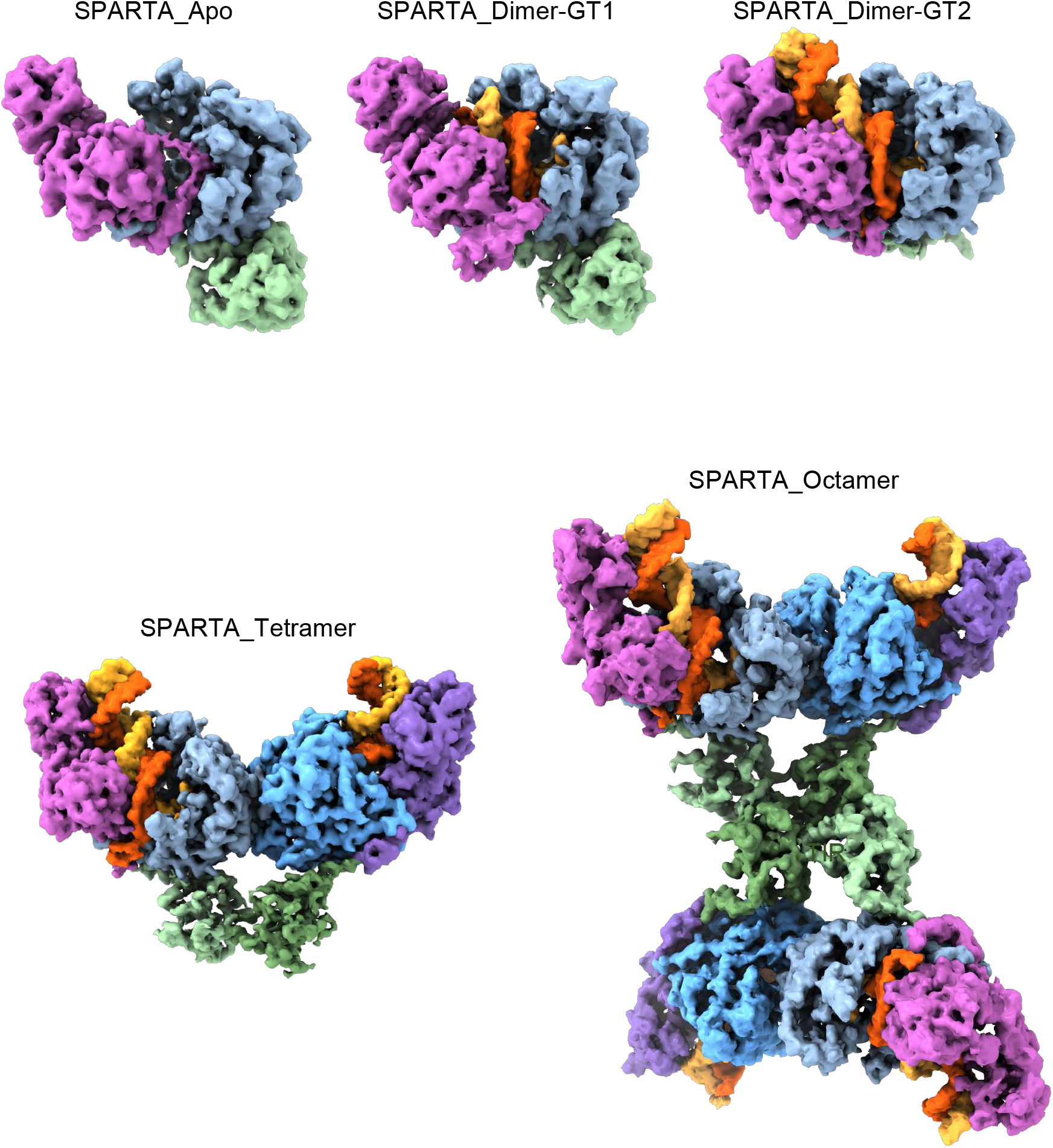
Summary of CrtSPARTA structures determined in this study. The 3D Cryo-EM maps of each structure are shown.

**Extended Data Fig 3.**
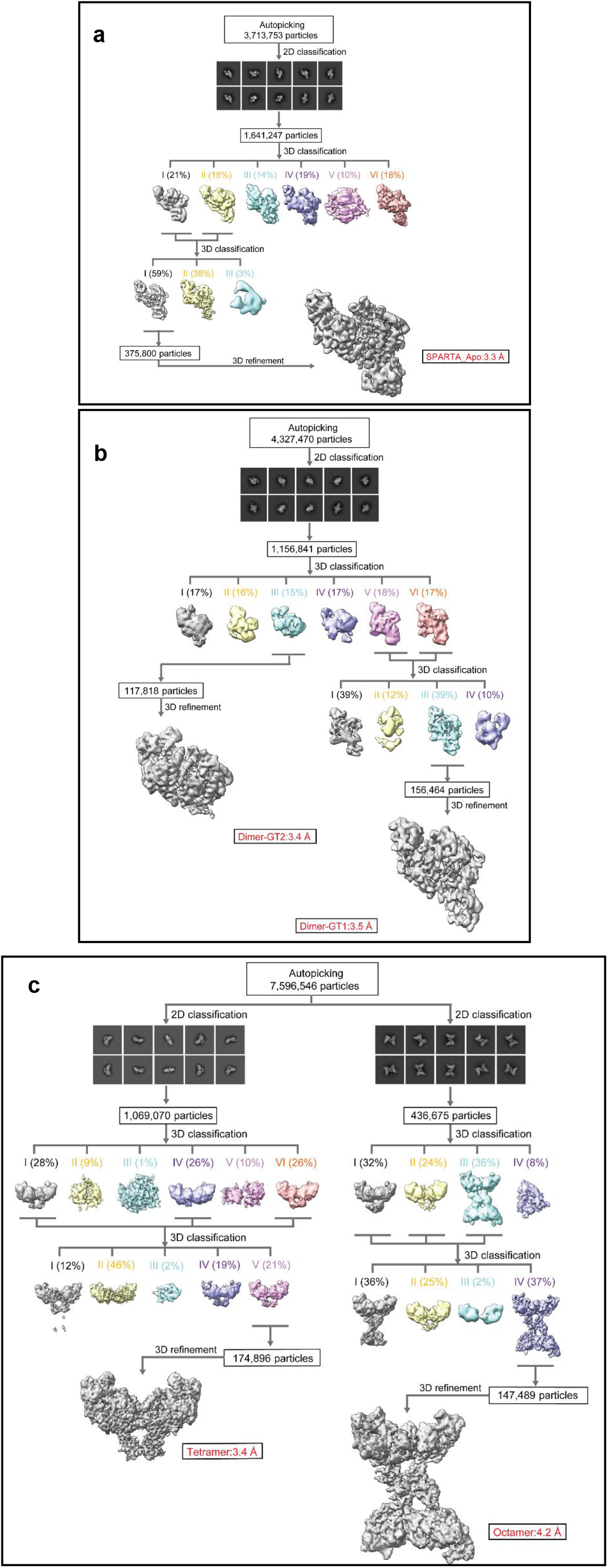
Single-particle cryo-EM analysis of CrtSPART apo (a), CrtSPARTA heterodimer bound with guide-target (b) and CrtSPARTA octamer sample (c).

**Extended Data Fig 4.**
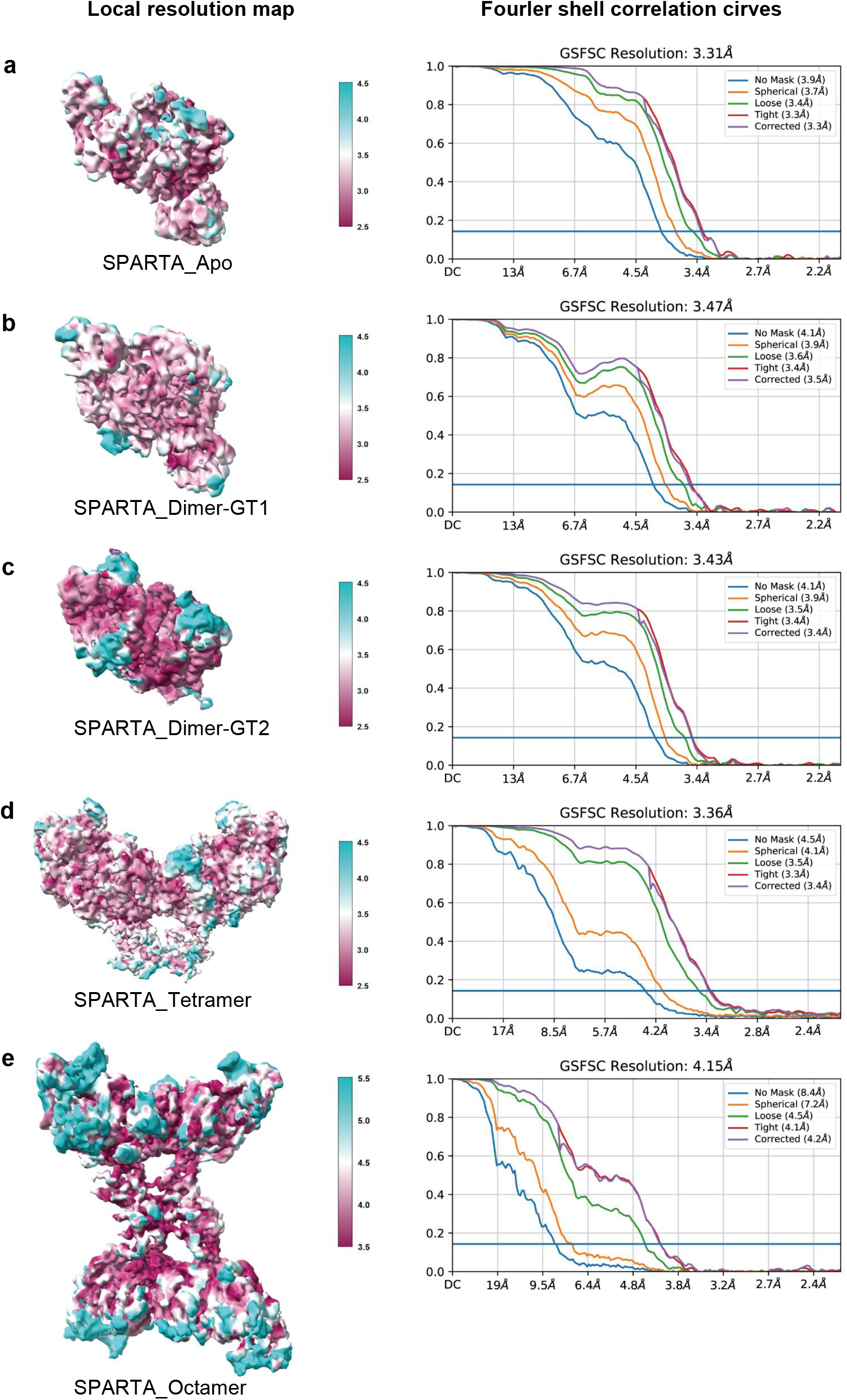
Cryo-EM density maps of the CrtSPARTA complexes. Maps are colored by local resolution, and gold-standard FSC of 0.143 resolution graphs is indicated for each complex.

**Extended Data Fig 5.**
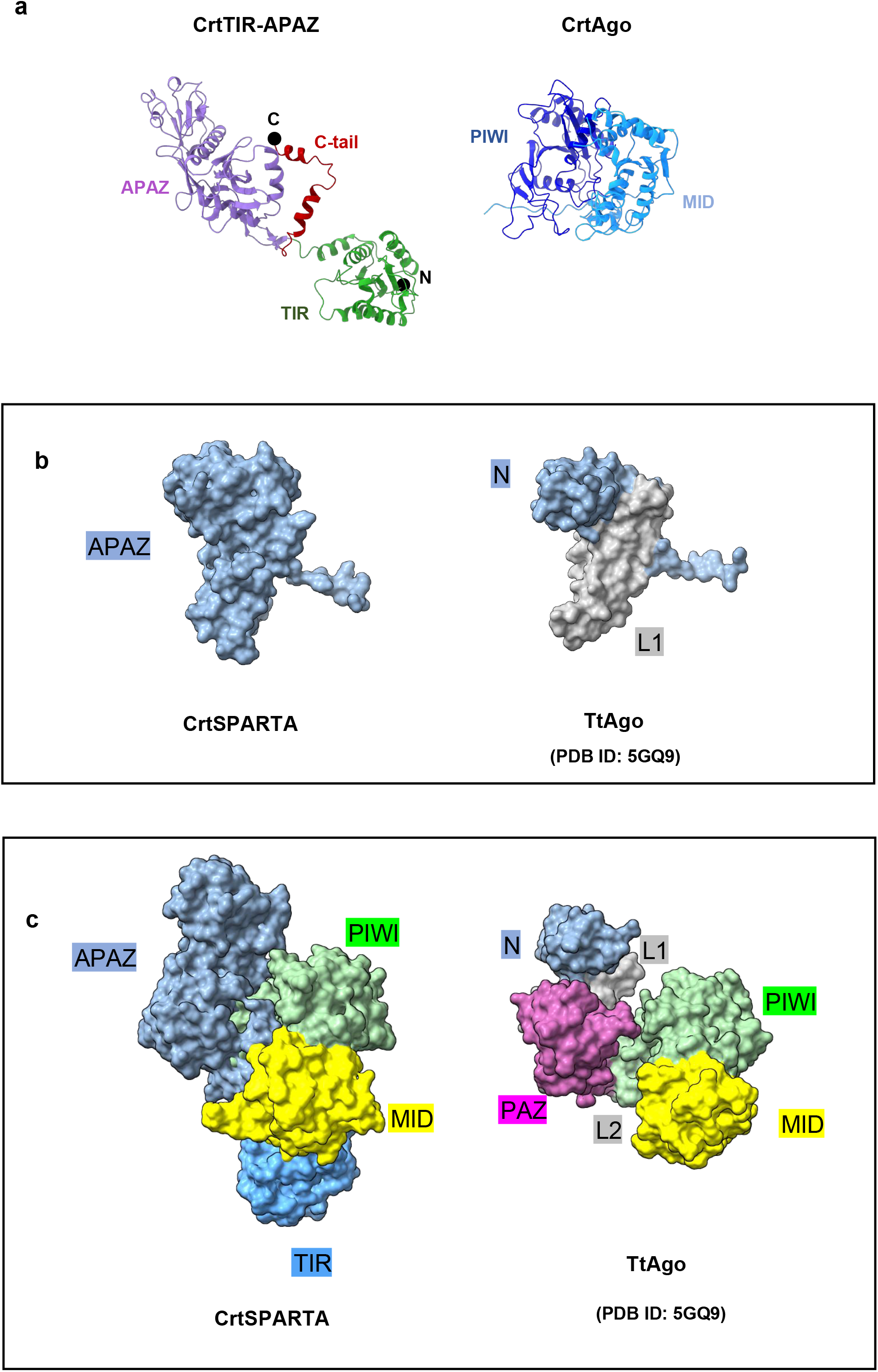
The structure of CrtSPARTA apo form. **a.** CrtTIR-APAZ and CrtAgo are shown side-by-side with each domain highlighted. **b.** The surface representation of APAZ and N domain connected with L1 linker from TtAgo (PDB:5GQ9). **c.** The overall structure comparison of CrtSPARTA with TtAgo (PDB:5GQ9).

**Extended Data Fig 6.**
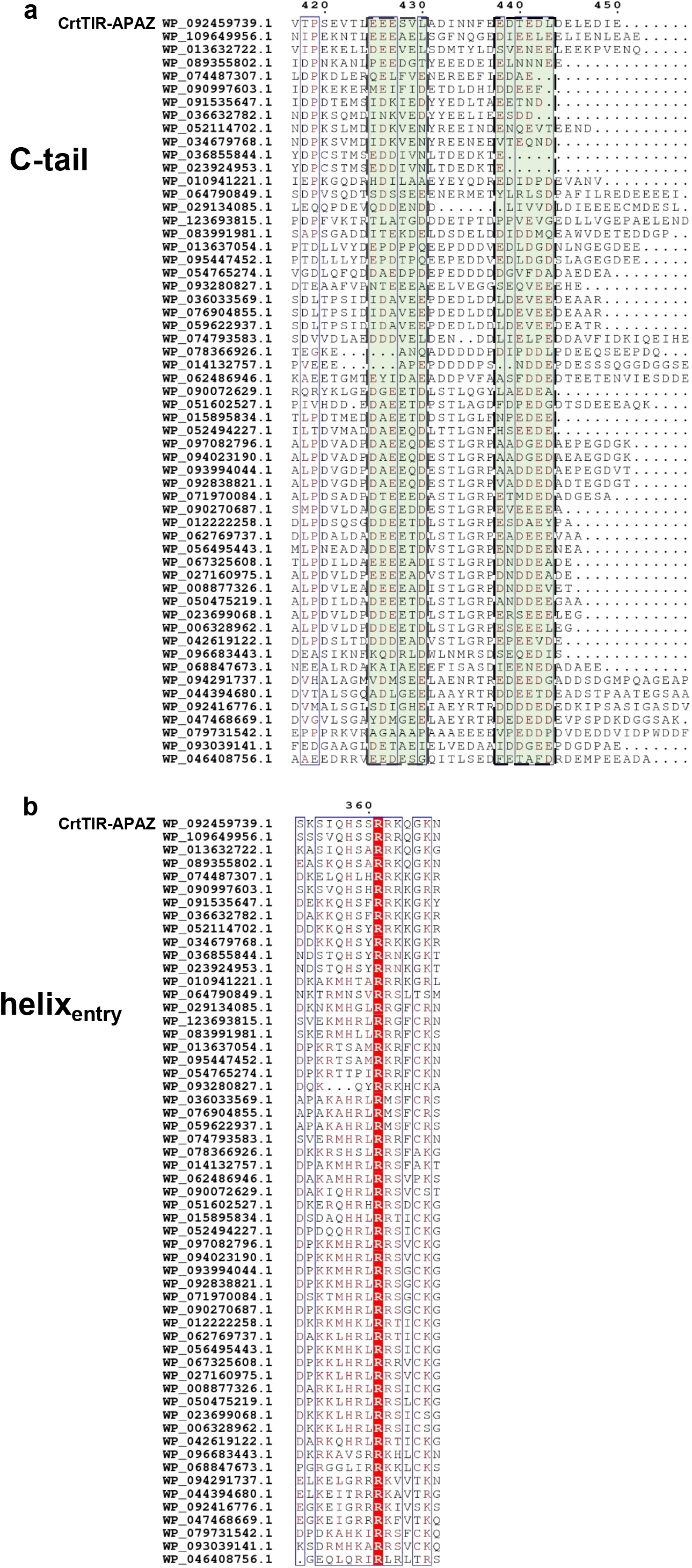
Sequence alignment of TIR-APAZ from 56 SPARTA systems. a. The sequence alignment of C-tail region. b. Sequence alignment of helixentry region. The GeneBank accession code for TIR-APAZ of each SPARTA system is listed in front.

**Extended Data Fig 7.**
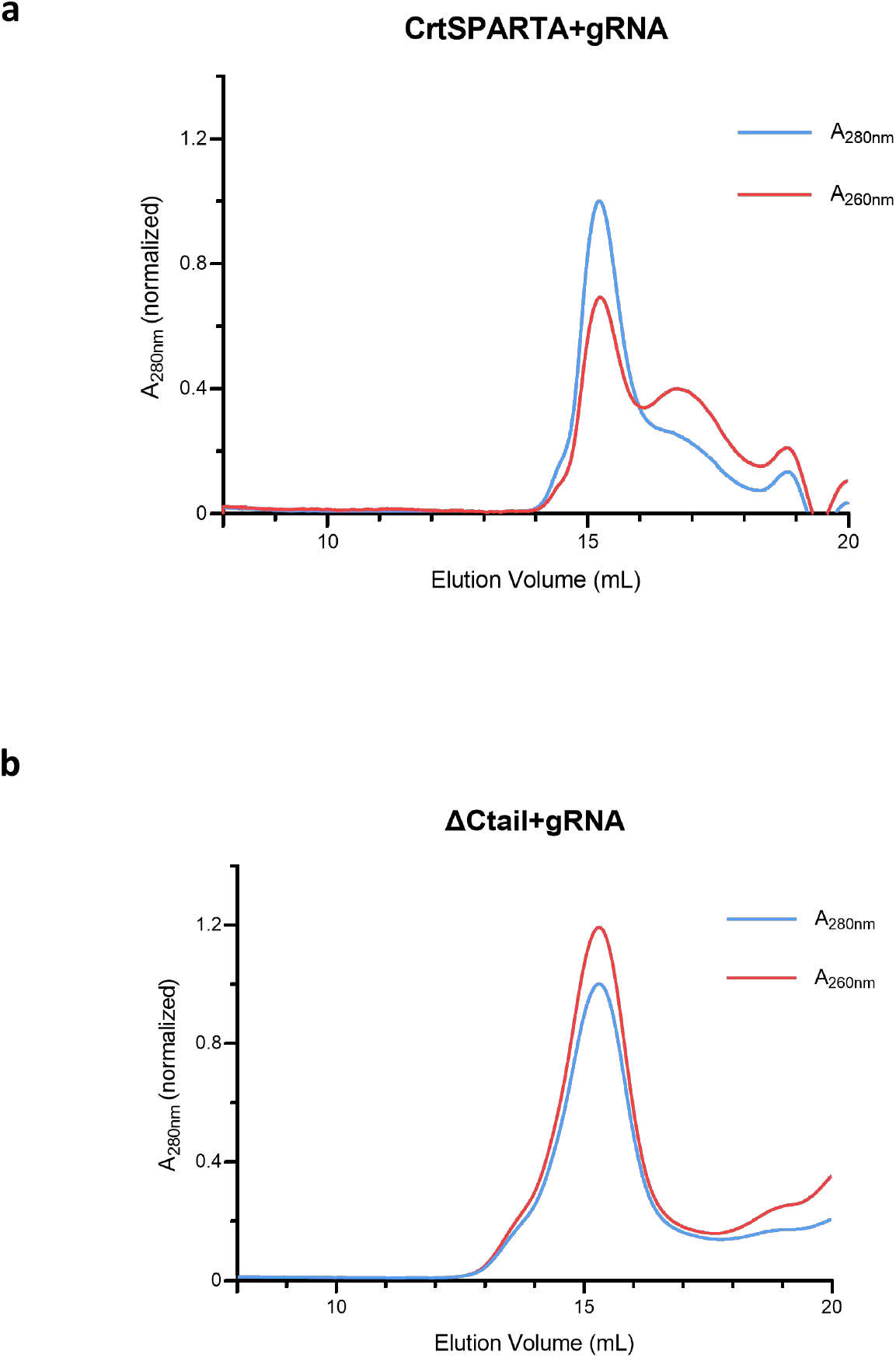
The size exclusion chromatography of (a) wild-type CrtSPARTA and (b) C-tail truncated CrtSPARTA (SPARTA_ΔCtail) incubated with guide RNA. The absorption at 260 nm and 280 nm is monitored and shown in red and blue, respectively.

**Extended Data Fig 8.**
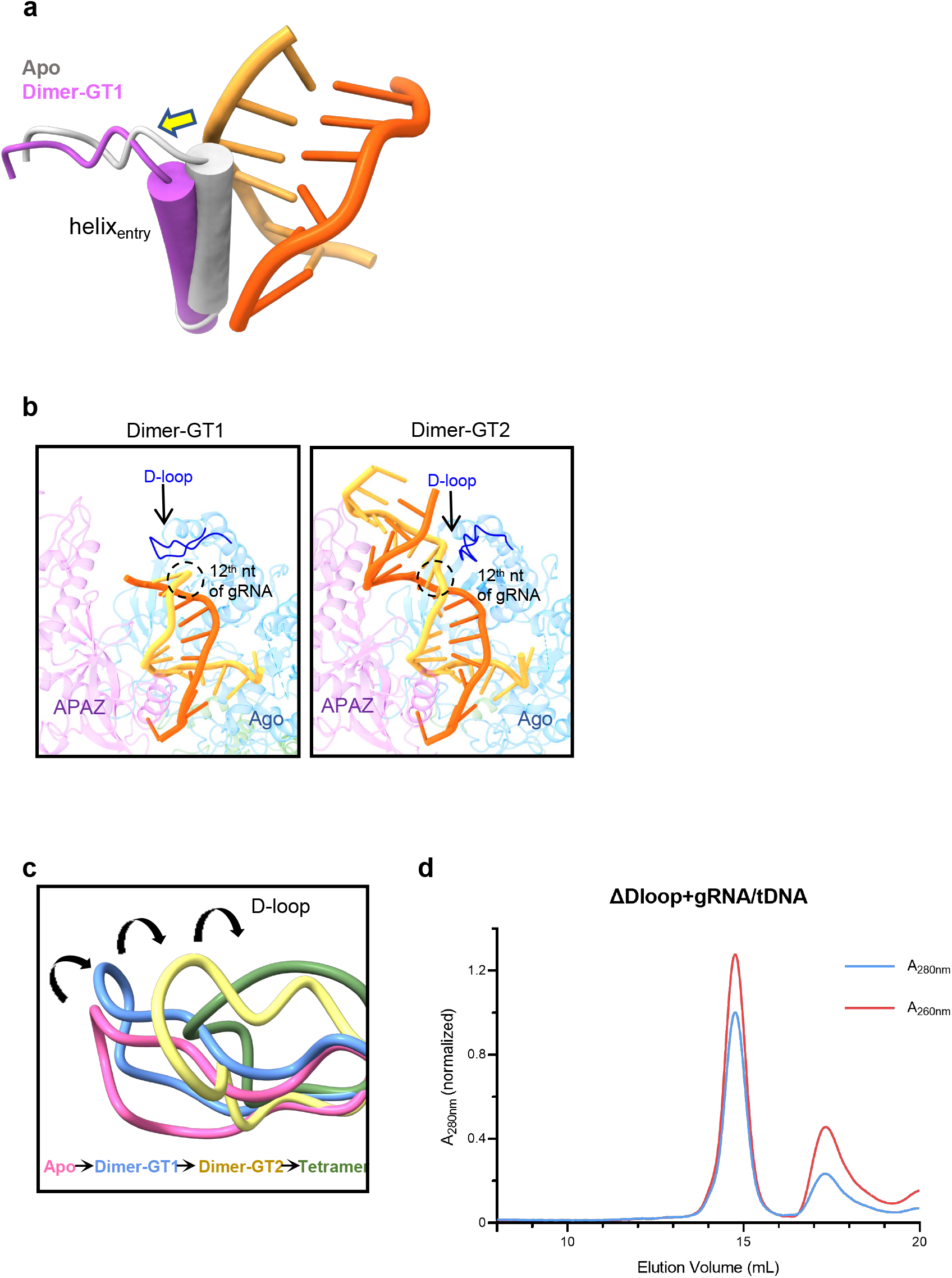
Guide-target binding induces the withdrawal of helix_entry_ and the flipping of the D-loop. a. The cartoon representation of helix_entry_ withdrawal upon guide-target binding. b. D-loop flipping along the propagation of guide-target. c. Structure superimposition of CrtSPARTA apo, Dimer-GT1, Dimer-GT2 and tetramer shows consecutive and unidirectional movement of the D-loop. d. The size exclusion chromatography shows D-loop truncated SPARTA cannot form octamer.

**Extended Data Fig 9.**
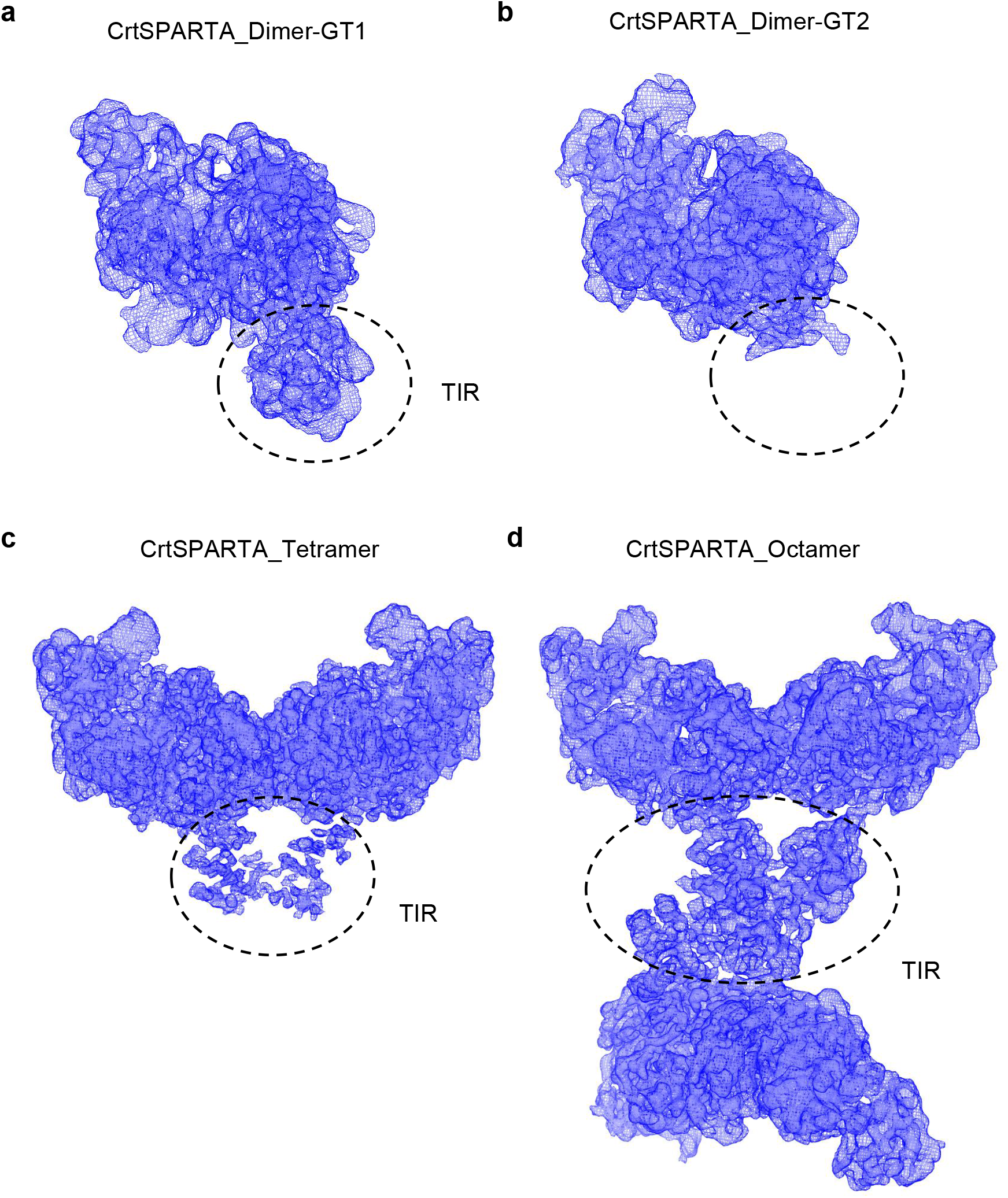
The Cryo-EM maps showing the flexibility of TIR domain. Comparison of TIR density in Dimer-GT1 (a), Dimer-GT2 (b), tetramer (c), and octamer (d).

**Extended Data Fig 10.**
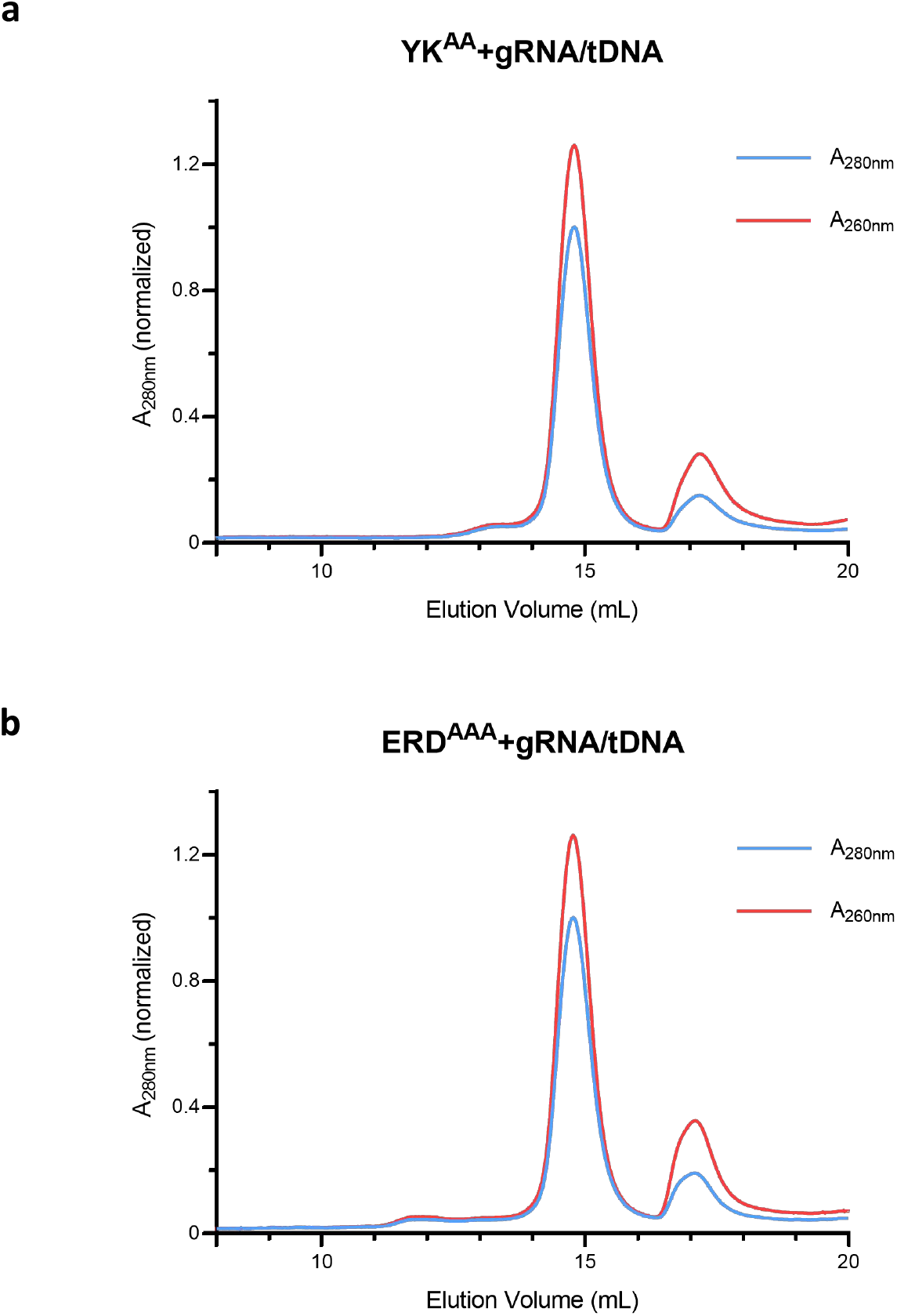
The size exclusion chromatography of the mutant CrtSPARTA_YK^AA^ (a) and CrtSPARTA_ERD^AAA^ (b) incubated with guide RNA and target DNA. The absorption at 260 nm and 280 nm are monitored and shown in red and blue respectively.

**Extended Data Fig 11.**
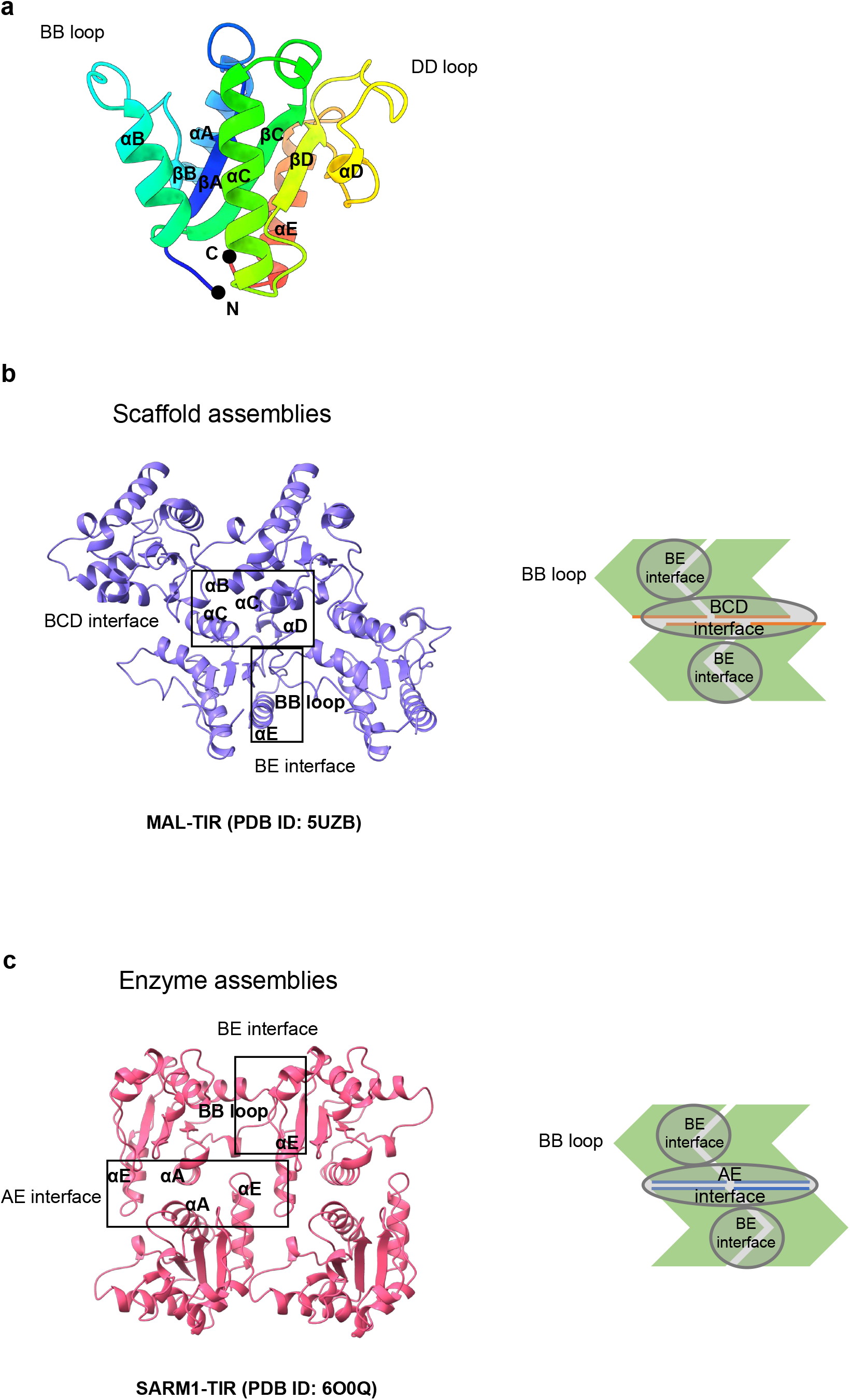
The typical assemblies of TIR domains. a. The structure of TIR domain in CrtSPARTA. The secondary structures are indicated. Eukaryotic TIR domains are primarily arranged either as “scaffold assembly” (b) or “enzymatic assembly” (c), corresponding to parallel or anti-parallel two stranded assembly via distinct interfaces, respectively. The involved interfaces are indicated by the frames.

**Extended Data Table 1.**
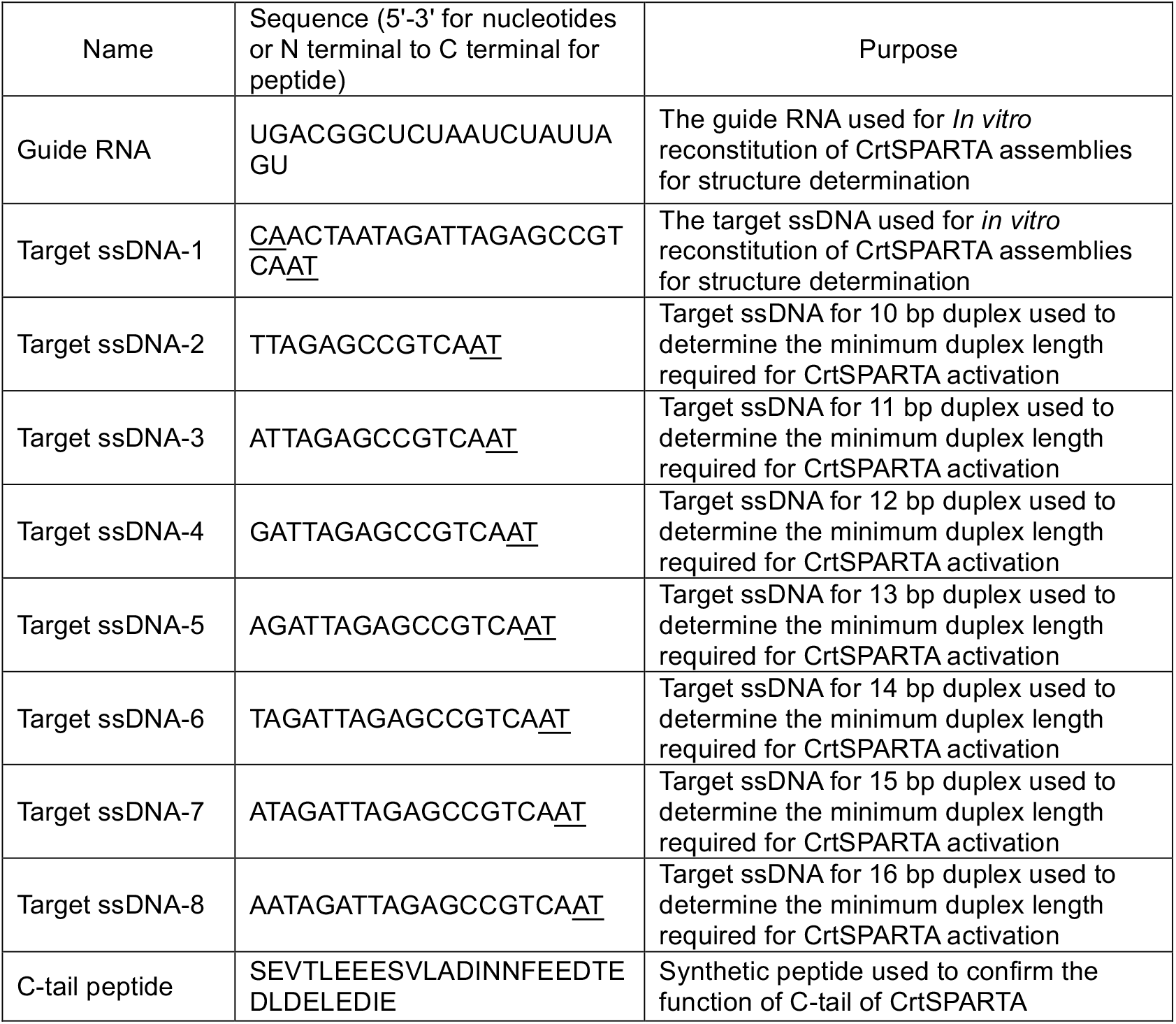
Sequences of nucleotides and synthetic peptide used in the study.

**Extended Data Table 2.**
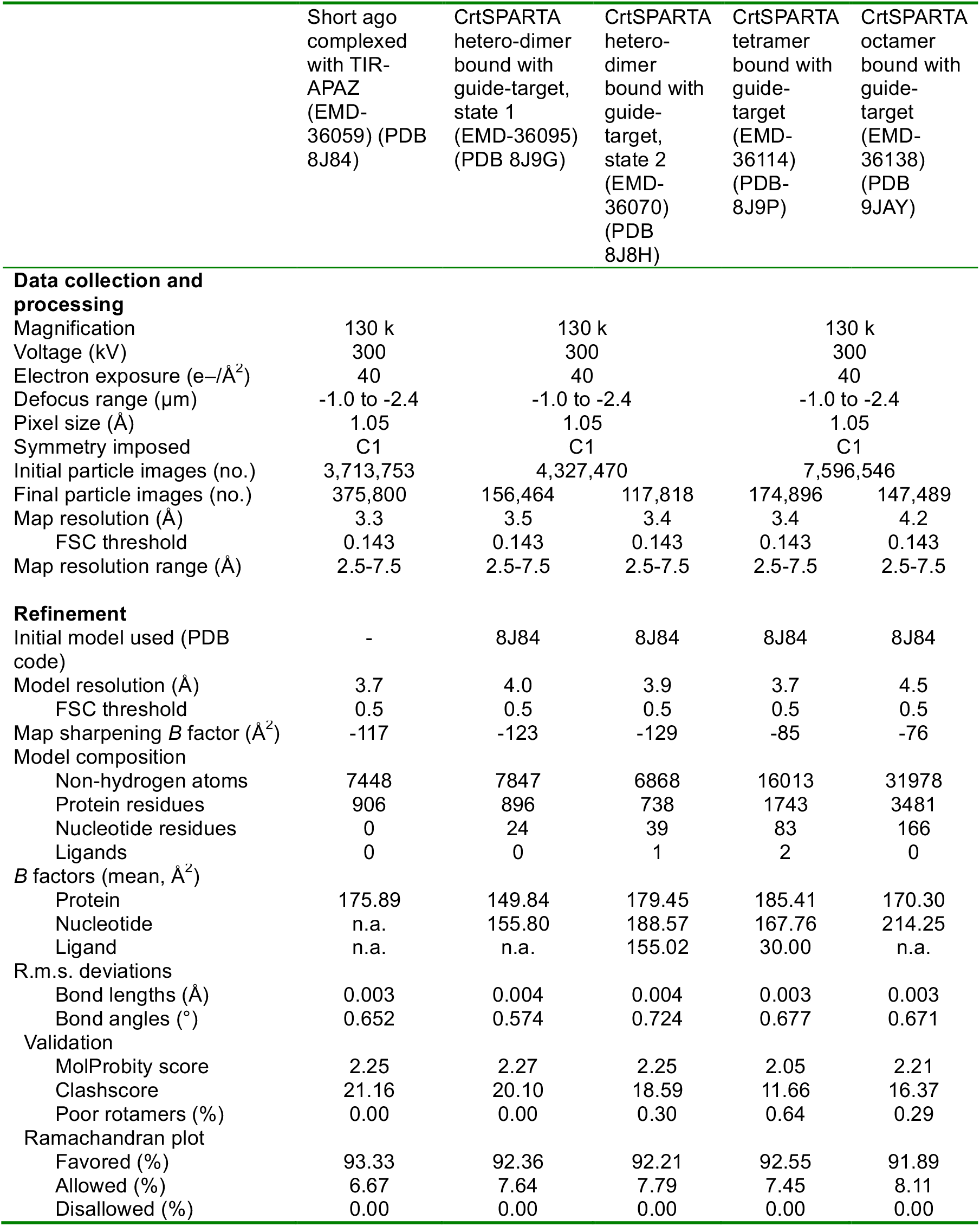
Cryo-EM data collection, refinement and validation statistics.

